# Developmental expression of the skeletal muscle determination gene, *MyoD*, is regulated by novel enhancer elements that interact with the core enhancer and distal regulatory region

**DOI:** 10.64898/2026.07.29.741533

**Authors:** Heather K. Jamieson, James R. Camp, Katherine Fleck, Cory J. Jubinville, Erica Korolev, Jennifer C. J. Chen, Leighton J. Core, Jelena Erceg, Masakazu Yamamoto, David J. Goldhamer

## Abstract

*MyoD* plays a central role in determining the skeletal muscle lineage in vertebrate embryos. The core enhancer (CE) and distal regulatory region (DRR) are the only known *MyoD* enhancers, and together, they recapitulate all major aspects of *MyoD* expression in the embryo. However, knocking out each enhancer individually has only modest effects on *MyoD* expression. Here, we show that embryos lacking both enhancers maintain muscle-specific *MyoD* expression, indicating the existence of unknown *MyoD* regulatory elements. Precision run-on sequencing together with available ChIP-seq and DNase I hypersensitivity datasets identified three new candidate enhancer regions within 96 kb of *MyoD* 5’ flanking sequences. Transgenic analysis revealed that DNA elements at -36 and -60 kb are active in all muscle-forming regions, each recapitulating aspects of endogenous *MyoD* expression. Enhancer activities in muscle regulatory factor-deficient mice suggest that they are components of the auto- and cross-regulatory circuitry that maintains *MyoD* expression. Analysis of Hi-C data showed that the entire -96 kb region constitutes a loop domain, within which multiple interactions between elements and with the *MyoD* gene were detected. A larger loop domain delimited by CTCF sites was also identified from -96 kb to +220 kb relative to the *MyoD* transcriptional start site. These data indicate that the newly identified enhancers are key components of a *cis* regulatory network that controls the activation and maintenance of *MyoD* expression in the embryo.

**One sentence summary:** *Cis* regulation of *MyoD* transcription during development

## INTRODUCTION

Members of the *MyoD* family of muscle regulatory genes, which include *MyoD*, *Myf5*, *Mrf4*, and myogenin, serve as nodal points in complex signaling and gene regulatory networks that control skeletal myogenesis in the embryo (Lima and Relaix, 2021; Weintraub et al., 1991). Collectively, the *MyoD* family is essential for myogenesis, although knockout (KO) studies by many laboratories have demonstrated partial genetic redundancy within this gene family (Hernández-Hernández et al., 2017). Each family member exhibits a distinct spatiotemporal pattern of expression in mouse embryos (Buckingham, 1992) and the extent to which gene-specific KO phenotypes reflect unique functional properties or expression patterns has not been fully established (Conerly et al., 2016; Gensch et al., 2008; Haldar et al., 2008; Tallquist et al., 2000; Wood et al., 2013). Nevertheless, it is known that *MyoD*, *Myf5*, and *Mrf4* function in establishing myoblast identity, with *MyoD* and *Myf5* being the primary drivers of lineage determination (Kablar et al., 1999, 1998, 1997; Kassar-Duchossoy et al., 2004; Rudnicki et al., 1993; Tajbakhsh et al., 1996). Myogenin is an essential regulator of muscle differentiation (Hasty et al., 1993; Nabeshima et al., 1993; Rawls et al., 1995; Venuti et al., 1995), which also requires either *MyoD* or *Mrf4* (Rawls et al., 1998).

Understanding muscle determination and differentiation pathways requires knowledge of how embryonic signals are integrated at DNA enhancer elements that control the appropriate activation and maintenance of muscle regulatory gene expression. Our previous work using transgenic mouse models indicated that DNA regulatory information contained within 24 kb of human *MyoD* 5’ flanking sequences (corresponding to approximately 27.5 kb of upstream sequence in the mouse) is sufficient to accurately control *MyoD* expression in mouse embryos (Chen et al., 2001). Activity of this genomic region could be accounted for by the combined activity of a 4-kb region centered at approximately -20 kb (designated fragment 3) (Chen et al., 2001; Faerman et al., 1995; Goldhamer et al., 1995, 1992) and the distal regulatory region (DRR) at -5 kb (Asakura et al., 1995; Tapscott et al., 1992) relative to the *MyoD* transcriptional start site (TSS). Molecular dissection and functional analysis of fragment 3 identified the 258-bp core enhancer (CE) at -20 kb (-23 kb in the mouse) which appears to be primarily responsible for the activity of the larger element (Chen et al., 2001; Faerman et al., 1995; Goldhamer et al., 1995). Analyses of enhancer-driven reporter gene expression in transgenic mice indicated that the CE plays a primary role in initiating *MyoD* expression, whereas the DRR functions to maintain and amplify *MyoD* expression levels (Chen et al., 2001; Faerman et al., 1995; Goldhamer et al., 1995; Kablar et al., 1999).

Regulatory mechanisms that control human and mouse *MyoD* expression through the CE and DRR have been extensively studied. A systematic linker-scanner mutagenesis screen of the human CE in transgenic mice identified multiple sequence elements that are essential for CE activity, including DNA elements specifically required for *MyoD* expression in myotomally-derived muscles (Kucharczuk et al., 1999). Available evidence is consistent with the notion that *MyoD* activation in the mouse is regulated by both the DNA and histone methylation status of the CE (Brunk et al., 1996; Scionti et al., 2017). Additionally, reporter gene assays in transgenic mice have shown that both the CE and DRR are targets of muscle regulatory factor (MRF)-dependent *MyoD* activation, with activity of the enhancers being either partially (CE) or completely (DRR) dependent on the MRFs (Kablar et al., 1999). More recent studies have shown that transcription of the mouse CE, which is regulated by the histone demethylase, LSD1 (Scionti et al., 2017), produces a non-coding enhancer RNA (^CE^RNA) that positively regulates *MyoD* transcription (Mousavi et al., 2013). The mouse DRR also encodes an enhancer RNA, known as ^DRR^RNA (Mousavi et al., 2013) or MUNC (Mueller et al., 2015), which primarily functions as a positive regulator of a subset of MYOD target genes in *trans* (Cichewicz et al., 2018; Mousavi et al., 2013; Mueller et al., 2015; Tsai et al., 2018), but also facilitates *MyoD* transcription (Mueller et al., 2015).

Given these studies, it is perhaps surprising that knocking out either the CE or DRR has only modest effects on *MyoD* expression in the embryo. Overall *MyoD* transcript abundance was reduced by approximately 30% in embryonic day 11.5 (E11.5) and E15.5 embryos lacking the CE (Chen and Goldhamer, 2004). Consistent with a role in initiating *MyoD* expression, CE KO embryos exhibited a delay of 1-2 days in expression of *MyoD* in the hypaxial migratory lineages that give rise to branchial arch and limb bud musculature (Chen and Goldhamer, 2004). The compensatory mechanisms that restore normal *MyoD* expression by E11.5 remain unknown. Further, it is well-established that in *Myf5* KO mice in which expression of the linked *Mrf4* gene is also abrogated (Kassar-Duchossoy et al., 2004), *MyoD* expression in trunk musculature is dependent on *Pax3* (Tajbakhsh et al., 1997). However, while a region within the CE was identified that is required for *Pax3*-dependent enhancer activity in transgenic mice (Chen and Goldhamer, 2004), the endogenous CE is dispensable for *Pax3* and *Myf5*-dependent regulation of *MyoD* (Chen and Goldhamer, 2004). In DRR KO mice, qualitative assessment by in situ hybridization suggested a modest reduction in *MyoD* transcript levels at E10.5, but quantitative analysis showed control levels of *MyoD* transcripts by E11.5 (Chen et al., 2002). It has remained unclear whether the mild KO phenotypes represent partial functional redundancy between the CE and DRR or the existence of unknown enhancers that cooperate with the CE and DRR to direct appropriate *MyoD* expression.

By sequentially targeting the CE and DRR in mouse embryonic stem cells (mESCs), we have produced mice lacking both enhancers. Transcript levels were reduced in double enhancer KO (dKO) embryos, but *MyoD* continued to be expressed in a muscle-specific pattern, supporting the existence of unknown *cis*-acting *MyoD* regulatory elements. Using precision run-on sequencing (PRO-seq) (Mahat et al., 2016) in conjunction with analysis of available ChIP-seq and DNase I hypersensitivity datasets, three novel candidate enhancer regions were identified within 96 kb upstream of the *MyoD* TSS. Two putative enhancer regions exhibited muscle-specific activity in transgenic mice and showed enhancer-specific MRF dependencies. Analysis of Hi-C data derived from cultured myoblasts showed that the novel regions interact with each other, with the CE and DRR, and with the *MyoD* gene. Hi-C analysis also identified additional interaction sites downstream of *MyoD* that may represent uncharacterized regulatory regions. The known, newly identified, and putative enhancer elements reside within a chromatin loop domain extending from a putative CTCF insulator site at -96 kb to the *MyoD* gene, and a second, larger loop delimited by the -96 kb CTCF site and another at +220 kb relative to the *MyoD* TSS. These data suggest that *MyoD* transcription is regulated by at least four enhancer elements that interact in complex ways to regulate *MyoD* expression in the embryo.

## MATERIALS AND METHODS

### MyoD CE targeting vector

A PGKneo cassette flanked by loxP sites was excised from the plasmid, ploxPneo-1 (provided by Dr. Marisa Bartolomei), by digestion with EcoRI (5’ end) and XhoI (3’ end) and the resulting overhangs were filled in with Klenow (New England Biolabs Cat # M0212S). This fragment was inserted and ligated in reverse orientation between two FRT sites in the loxP2-FRT2/BSIIPSK plasmid that had been blunt end-linearized with EcoRV.

The FRT-loxP-PGKneo-loxP-FRT (FLneoLF) portion of the plasmid was removed via digestion with NotI and EcoRI and inserted into p5’enh3’EB (Chen and Goldhamer, 2004), which contains the CE with 3.7 kb and 3.6 kb of genomic DNA 5’ and 3’ of the CE, respectively. The CE-containing fragment was excised by digestion with EcoRI and BamHI, which had been introduced at the 5’ and 3’ ends of the CE, respectively (Chen and Goldhamer, 2004), and the FLneoLF fragment inserted in its place. The resulting ligation product was designated p5’FLneoLF3’. A thymidine kinase (tk) cassette was added to the 3’ end of p5’FLneoLF3’ at the XhoI site. The plasmid, p5’FLneoLF3’tk, was grown in NM544 electrocompetent bacteria and the plasmid was isolated using Qiagen’s EndoFree Maxi Kit, as per the manufacturer’s instructions.

### Production of CE/DRR dKO mice

Given the close linkage of the CE and DRR on Chromosome 7, mESCs lacking both the CE and DRR were produced by sequential targeting of the enhancers. UCONN Health’s Center for Mouse Genome Modification (CMGM) produced mESCs derived from blastocysts produced by crossing DRR KO mice (Chen et al., 2002), in which the *MyoD* genomic region was derived from the 129SvJ strain, with 129SvEv mice. The CE targeting vector’s homology arms were derived from the 129SvJ strain, with the intended purpose of increasing the efficiency of targeting the 129SvJ chromosome, which lacked the DRR. The CE targeting vector was linearized with NotI, and the purified fragment resuspended in TE at a concentration of 1 μg/μl. Electroporation of the CE targeting vector and isolation of mESC clones was performed by the CMGM.

mESC clones were screened for proper recombination by Southern blotting on the 5’ end and nested long-range PCR on the 3’ end. The Southern probe was PCR amplified from mouse genomic DNA using the forward primer 5’-GGCGGATCCTGAACAAAAGGGGATGAGATTCC-3’ and reverse primer 5’-CGCGAATTCAGGAACCACCCTAAAGATCCACC-3’ and subcloned into pBS SK+. The Invitrogen Elongase system (Cat # 10480-010) was used for long-range PCR. The external PCR primer sequences were 5ʹ-AGTAGAAGGTGGCGCGAAGG-3’ and 5’-TCAAGCCGGCCACCATAAAG-3’. The internal PCR primers were 5’-TCATTCAGGAGAGCCTTTGTT-3’ and 5’-TGGATGTGGAATGTGTGCGAG-3’. The final amplicon was 4.6 kb.

Chimeric mice were produced by the CMGM using standard methods, and highly chimeric mice were mated with FVB mice, and offspring genotyped for both the CE^neo^ and DRR^loxP^ alleles. CE^neo/+^;DRR^loxP/+^ male mice were crossed with females harboring the *R26^Flpe^* allele (Jax stock #003946; (Farley et al., 2000) to remove the neo cassette from the CE locus (designated *MyoD^ΔCD/+^* mice), or with *Hprt^Cre^* females (Jax stock #004302; (Tang et al., 2002) to generate mice in which the neo cassette and all sequences between the CE and DRR were removed (designated *MyoD^ΔC18D/+^*mice; Fig. 1). Double enhancer KO alleles were detected by PCR, and primers used and corresponding amplicon sizes are provided in Supplemental Table 1.

**Figure 1.**
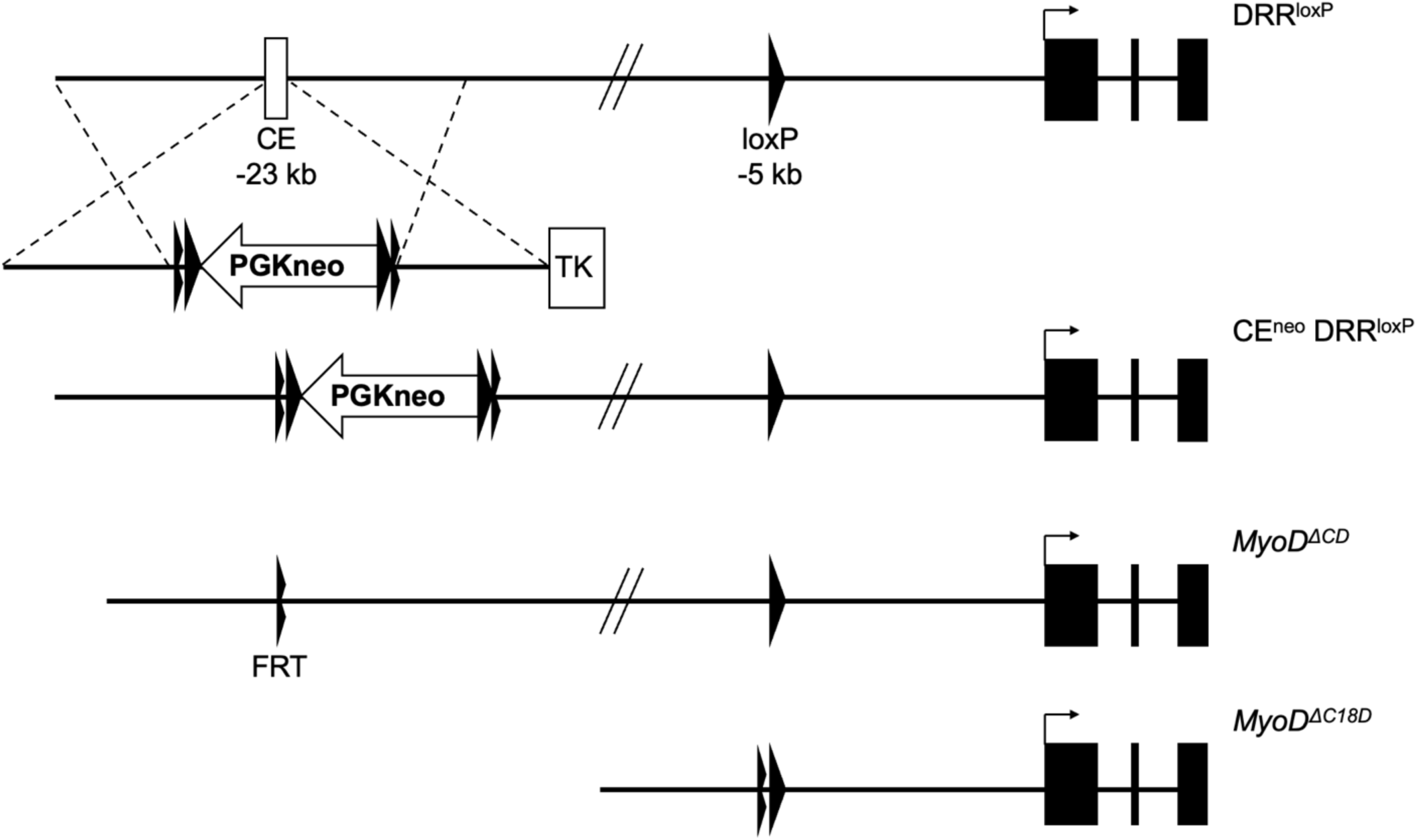
Sequential targeting strategy to produce double enhancer knockout lines. The core enhancer (CE) was replaced with a PGKneo cassette in ESCs derived from DRR^loxP^ mice, in which a single loxP site replaced the distal regulatory region (DRR). PGKneo is flanked by both flippase recognition target sites (FRT) and loxP sites. Flippase-mediated recombination yielded *MyoD^ΔCD^* lines, in which the CE and DRR are replaced with FRT and loxP sites, respectively. Cre-mediated recombination generated *MyoD^ΔC18D^* lines, which lack both enhancers, as well as ∼18 kb of genomic sequence between them.

### MyoD quantification of enhancer KO mice

Collected embryos were kept on ice until being processed. The limb buds and trunk of E12.5 embryos were separately dissected and immediately lysed in Qiagen RLT lysis buffer supplemented with 10 µL of β-mercaptoethanol per 1 mL of lysis buffer. Samples were disrupted by pipetting and then homogenized with a Qiagen QIAshredder column. RNA was purified with a Qiagen RNeasy Mini column followed immediately by cDNA synthesis using the Bio-Rad iScript Advanced cDNA Synthesis kit. After cDNA synthesis, *MyoD* transcripts were quantified by ddPCR using a TaqMan probe and intron-spanning primers (Applied Biosystems, Catalog No: 4331182; Assay ID: Mm00440387_m1). Statistical comparisons used one-way ANOVA with Tukey’s multiple comparisons.

### Generation of MyoD candidate enhancer reporter constructs and transgenic lines

Three candidate *MyoD* enhancer regions, designated as the -96 kb, -60 kb, and -36 kb regions, were subcloned into the -2.5lacZ plasmid (Goldhamer et al., 1992) using standard recombineering techniques (Liu et al., 2003). The size of each fragment and its relationship to discriminative regulatory element detection from GRO-seq (dREG) (Danko et al., 2015) regions identified by PRO-seq (see below) is provided in Supplemental Table 2. For each candidate region, two PCR products that subsequently served as homology arms were amplified via PCR with the following primer sets. -96 kb 5’ arm: 5’-GCTGTCGACGCCAACAGTGACAACCGAAG-3’ and 5’-GTTTTCGAACCACACAAGGAGCGAGTTCT-3’, -96 kb 3’ arm: 5’-TTGTTCGAACCTGCGAGCACATGTGTAGA-3’ and 5’-GTGACTAGTT CAACCACGCTCAACTGCAT-3’; -60 kb 5’ arm: 5’-GGAGTCGACAGTCCTGTGTGTGCATGTGA-3’ and 5’-CTGTTCGAATCCTCCCAAGACTCCATCCA-3’, -60 kb 3’ arm: 5’-TAGTTCGAAGAGAAAGGTAGACAGAAGGCCT-3’ and 5’-AGAACTAGTGGCTCTGCTTGGATGTCTCT-3’; -36 kb 5’ arm: 5’-AAAGTC GACCTAGGAATCTCACAGAACTGC-3’ and 5’-GCTTTCGAATCCACCTGCCTAGACAAGAA-3’, -36 kb 3’ arm: 5’-TTATTCGAAGGTTGTGAGCCCAGACATGA-3’ and 5’-TCAACTAGTAGGAAACACAGGCAG CATCA-3’. 5’ and 3’ PCR products were digested with SalI/BstBI and BstBI/SpeI, respectively, and inserted together into the SalI/SpeI-digested -2.5lacZ plasmid by a three-fragment ligation, resulting in retrieval plasmids. To generate -96lacZ and -36lacZ transgene constructs, gap-repair retrieval of the intervening sequence was performed via recombineering with BstBI-digested retrieval plasmid, RP23-65K18 BAC DNA (Invitrogen), and EL350 host bacteria, as described previously (Liu et al., 2003). Although the -60 kb candidate region was retrieved via recombineering, the generated plasmid was not stable. Re-cloning of the blunted BsrGI-SpeI fragment of the retrieved -60 kb candidate region into the blunted SpeI site of -2.5lacZ resulted in a stable -60lacZ transgene construct. The transgene plasmids were prepared using the QIAGEN Extra-Midi Plasmid Kit. -96lacZ and -36lacZ transgene cassettes were excised from the vector backbone by NotI/SalI digestion. The -60lacZ transgene was isolated by NotI digestion. Purification of excised transgene cassettes and pronuclear injection into C57BL/6J one-cell embryos were performed by the CMGM.

### Mouse maintenance, breeding, and genotyping

All mouse procedures were approved by the University of Connecticut IACUC. Mice were maintained on an enriched FVB background. Standard crossing schemes were used to generate experimental animals, and specific crosses are presented in the Results. Mice were genotyped by PCR from DNA isolated from tail snips (adults) or embryo-derived extraembryonic membranes. PCR primers and corresponding amplicon sizes are provided in Supplemental Table 1.

### Embryo collection, whole-mount in situ hybridization, photography, and imaging

For all timed matings, noon on the day of the vaginal plug was considered E0.5. Embryos were collected at times specified in the Results and individually fixed in 4% PFA in PBS overnight at 4°C with gentle rocking. Mice were then processed for in situ hybridization as previously described (Yamamoto et al., 2007). Whole-mount images were captured with a Leica MZFLIII stereomicroscope (Spot RT3 digital camera) or with an Olympus SZX10 (Pentax K-30). Focus stacking, minor adjustments in brightness, midtone, color, and contrast were made in Adobe Photoshop, Affinity Photo 2, or GIMP.

### Satellite cell isolation and culturing

Cells were isolated via fluorescence-activated cell sorting (FACS) as described previously (Yamamoto et al., 2018), or via magnetic-activated cell sorting (MACS). For both FACS and MACS isolation, mice were sacrificed and muscles of the lower hindlimb were mechanically and enzymatically digested. For FACS isolation, cells were then stained, and CD31^-^/CD45^-^/α7-integrin^+^/VCAM^+^ cells were collected and plated in growth media (GM; 20% FBS [HyClone, Cat # SH30071.03], 2.5 ng/mL basic-FGF [Life Technologies, Cat # PHG0264], and 1x Pen/Strep in DMEM) on tissue culture plates coated with 1 mg/mL Matrigel (Corning, Cat # 356231).

For MACS isolation, the Satellite Cell Isolation Kit (Miltenyi, Cat # 130-104-268) was used, followed by positive selection with anti-Integrin α-7 microbeads (Miltenyi, Cat # 130-104-261). Cells were plated in GM, and either remained in GM or were switched into differentiation media (DM; 2% horse serum and 1x Pen/Strep in DMEM) when cells reached approximately 80% confluence. After 3 days in DM, cells were fixed in 2% PFA/0.25% glutaraldehyde in PBS for 10 min at 4°C and stained for β-galactosidase, as previously described (Yamamoto et al., 2007).

### Cell permeabilization for PRO-seq

All steps were performed on ice with ice-cold buffers. Satellite cells in 10 cm dishes were briefly washed twice in PBS, followed by scraping in 5 mL Buffer P (10 mM Tris-HCl pH 8.0, 10% glycerol, 10 mM KCl, 5 mM MgCl_2_, 250 mM sucrose, 0.5 mM DTT, 0.1% IGEPAL, 1 unit/mL RNase inhibitor (SUPERaseIN, Ambion)), and transferred to a 15 mL conical tube. The plates were rinsed and scraped with an additional 5 mL of Buffer P. Cells were pelleted by centrifugation at 800 x g for 5 min in a swinging bucket rotor. Cell pellets were washed twice by resuspension in 5 mL of Buffer W (Buffer P without IGEPAL) and centrifugation at 800 x g for 5 min. Permeabilized cells were resuspended in Buffer F (5 mM Tris-HCl pH 8.5, 40% glycerol, 5 mM MgCl_2_, 0.5 mM DTT, 5 mM DTT, 5 units/mL RNase inhibitor) at a density of ∼1 x 10^6^ cells/100 µL and immediately frozen in liquid nitrogen. Samples were stored at - 80°C until library preparation.

### PRO-seq library preparation

PRO-seq libraries were generated from permeabilized cells from biological replicates using the protocol described in Mahat et al. (2016), with the following modifications. 4-biotin run-on reactions were carried out in a final volume of 200 μL. After adding the run-on master mix, samples were vigorously pipetted for 30 seconds and then incubated at 37°C for 10 min. After the run-on reaction, RNA was extracted using Norgen RNA purification columns (Cat # 37500) as per the manufacturer’s protocol. Following purification, RNA was base-hydrolyzed for 20 min on ice prior to enrichment of biotinylated nascent RNA as in Mahat et al. (2016). 3’ RNA adapter ligation was carried out off-bead using 15 pmol adapter, followed by 5’ decapping, 5’ hydroxyl repair, and 5’ RNA adapter ligation performed on beads. Upon completion of reverse transcription, libraries were pre-amplified for 5 cycles using cycling parameters previously described (Mahat et al., 2016). Test amplifications using serial dilutions of the pre-amplified libraries were then performed to determine the ideal number of cycles for full-scale amplification, with 15 cycles chosen for all samples. Fully amplified libraries were purified using NEB Monarch PCR & DNA cleanup kits (Cat # T1030L), quantified by Qubit, pooled in an equimolar fashion, and submitted for single-end, 75bp sequencing on an Illumina NextSeq 500 at the Center for Genome Innovation (UConn, Storrs, CT). Libraries were sequenced to a total depth of ∼30 million uniquely aligned reads (mm10 genome), with alignments performed as in Mahat et al. (2016).

### Identification of transcription regulatory elements (TREs)

To identify regions of bidirectional transcription in PRO-seq data, strand-specific bigwig files representing the 3’ ends of reads were generated. First, replicate data was pooled and converted to strand-specific bedgraphs of 3’ ends using *bedtools genomecov*. Bedgraph files were then converted to bigwig format using the bedGraphToBigWig script from Kent Utils (https://www.encodeproject.org/software/bedgraphtobigwig/). The resulting bigwig files were used as input for the dREG *run_predict.bsh* script (https://github.com/Danko-Lab/dREG) (Danko et al., 2015). dREG regions were merged into ‘peaks’ of regulatory elements with the *writeBed.sh* script using a dREG score of 0.8 as a threshold.

### Hi-C and ChIP-seq data processing

BL-Hi-C raw sequencing reads for primary myoblasts at proliferation and early differentiation stages were obtained from CRA002490 (Wang et al., 2022)Supplemental Table 3). To generate Hi-C maps, the bridge linker was removed from the raw reads using trimLinker from ChIA-PET2 (Li et al., 2017), which were then processed according to the established 4D Nucleome (4DN) Hi-C Processing Pipeline (https://data.4dnucleome.org/resources/data-analysis/hi_c-processing-pipeline). Trimmed fastq reads were mapped to the mm10 genome assembly using bwa mem (v0.7.17) (Li, 2013), then filtered for duplicates and merged across replicates with pairtools (v0.3.0) (Open2C et al., 2024). Using a HaeIII restriction enzyme file, Hi-C maps were generated with Juicer Tools (v1.22.01) command juicebox-pre (Durand et al., 2016). Loops were annotated using HiCCUPS with parameters -r 5000 -k SCALE -f 0.1 - p 1 -i 3 -d 5000. In situ Hi-C data from mESCs was downloaded from 4DNFI3JYF9VS. SCALE normalization was added to this file using addNorm from Juicer Tools, then annotated for loops with HiCCUPS using the same parameters.

H3K27ac, H3K4me1, CTCF, and MyoD ChIP-seq raw sequencing reads were also obtained from CRA002490 (Wang et al., 2022) for both GM and DM conditions (Supplemental Table 3). ChIP-seq data was processed as previously described (Jusuf et al., 2026). Sequencing reads were mapped to mm10 using bowtie2 (v2.2.5) (Langmead and Salzberg, 2012), filtered to retain uniquely mapped reads with samtools (v1.21) (Danecek et al., 2021), duplicates removed with sambamba (v0.6.6) markdup, and merged across replicates with sambamba merge (Tarasov et al., 2015). Final bam files were converted to bedGraphs with bedtools (v2.29.0) bamtobed and genomecov (Quinlan and Hall, 2010). Processed bigWig files of H3K27ac, H3K4me1, and CTCF ChIP-seq data in mESCs were obtained through ENCODE (ENCFF583WVZ, ENCFF456IEO, and ENCFF144RYY). These files were converted to bedGraphs using bigWigToBedGraph from Kent tools (v1.04.00) (Kent et al., 2010).

For visualization, Hi-C and bedGraph files were directly loaded into the UCSC Genome Browser (Casper et al., 2025). The files produced by HiCCUPS were converted into Interact files before loading into the browser to visualize loop interactions as arcs.

## RESULTS

### Production of CE/DRR dKO mice

As knocking out the CE and DRR individually has only modest effects on *MyoD* expression (Chen et al., 2002; Chen and Goldhamer, 2004), we tested whether the CE and DRR serve partially redundant functions by producing mice that lacked both enhancers. Since the CE and DRR are closely linked on mouse Chromosome 7 (approximately 23 kb and 5 kb upstream of the *MyoD* TSS, respectively), CE/DRR dKO mice were produced by sequential targeting. Starting with mESCs in which the DRR on one chromosome was replaced with a loxP sequence, the CE on that chromosome was replaced with a neo cassette flanked by both FRT and loxP sites (CE^neo^ DRR^loxP^; Fig. 1). After generating heterozygous germline knockout mice, the neo cassette was removed by crossing these mice to *R26^FLPe^*mice (Farley et al., 2000) to generate mice lacking both enhancers on one chromosome. The resulting dKO allele is designated *MyoD^ΔCD^*. A second dKO allele that, in addition, lacks approximately 18 kb of sequence between the CE and DRR was produced by crossing CE^neo^ DRR^loxP^ mice to the Cre-deleter line, *Hprt^Cre^* (Tang et al., 2002; Fig. 1). This dKO allele was designated *MyoD^ΔC18D^*. Heterozygous lines were crossed to produce homozygous mice, which were viable and fertile. Both homozygous-null lines were produced at the expected Mendelian frequencies (data not shown).

### Muscle-specific expression of MyoD does not require the CE or DRR

We tested whether *MyoD* expression was more substantially affected in dKO embryos by quantifying *MyoD* transcript levels by ddPCR. This analysis was conducted on E12.5 embryos, when the pattern of *MyoD* expression in wild-type (WT) and single KO embryos is indistinguishable (Chen et al., 2002; Chen and Goldhamer, 2004). Limb buds and trunks were assayed separately to determine whether *MyoD* expression in migratory and myotomal populations is differentially affected. *MyoD* expression in *MyoD^ΔCD/ΔCD^* embryos was significantly reduced in the trunk (p = 0.034; 20.6% of WT) and trended lower in the limb buds but did not reach statistical significance (p = 0.079; 25.0% of WT; Supplemental Fig. 1).

Whole-mount in situ hybridization was performed on *MyoD^ΔCD/ΔCD^* and WT embryos from E9.75 to E12.5 to assess how the muscle-specific pattern of *MyoD* expression was affected by loss of both the CE and DRR (Fig. 2). In WT embryos, *MyoD* mRNA is detected as early as E9.5 in the somites and the first branchial arch (Fig. 2A) and by E10.5 and E11.5 in the forelimb and hindlimb buds, respectively (Fig. 2D, G) (Chen et al., 2002; Chen and Goldhamer, 2004). *MyoD* was expressed in *MyoD^ΔCD/ΔCD^* embryos, and expression approximated the additive effects of the individual enhancer knockouts. Like CE KO embryos (Chen and Goldhamer, 2004), branchial arch expression in *MyoD^ΔCD/ΔCD^* embryos was delayed but not abrogated (Fig. 2A, B, E). Staining of the forelimb buds of *MyoD^ΔCD/ΔCD^* embryos at E10.5 was reduced in intensity compared to WT embryos (Fig. 2D, E), approximately phenocopying *MyoD* expression in CE (Chen and Goldhamer, 2004) and DRR (Chen et al., 2002) KO embryos. No consistent expression differences were observed between *MyoD^ΔCD/ΔCD^* and *MyoD^ΔC18D/ΔC18D^*embryos (Fig. 2), indicating that the 18 kb of sequence between the CE and DRR does not serve a regulatory function in controlling *MyoD* expression. Collectively, these data indicate that the CE and DRR do not serve redundant functions in regulating the muscle specificity of *MyoD* expression and point to the existence of additional *MyoD* regulatory elements.

**Figure 2.**
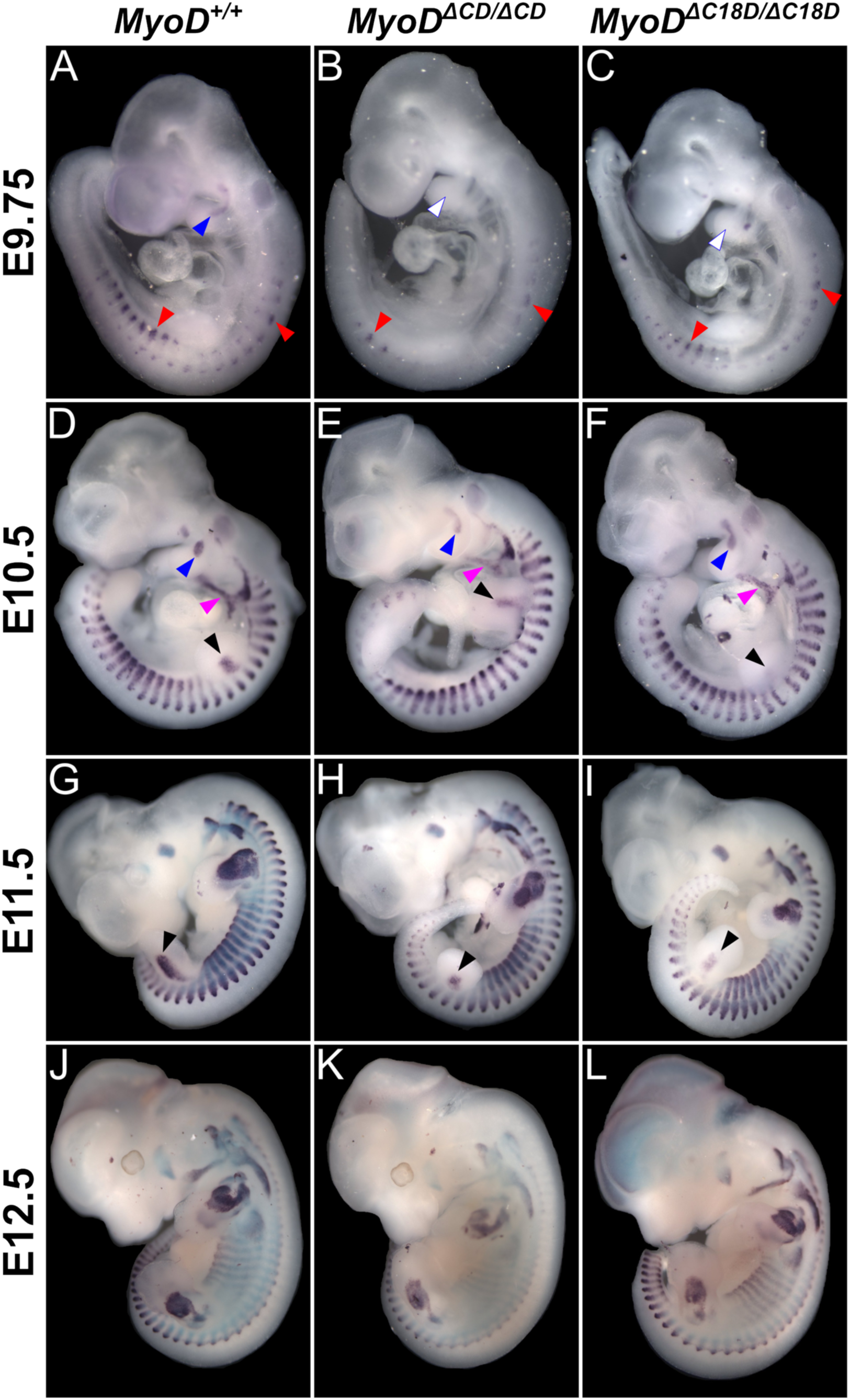
Persistent muscle-specific expression of *MyoD* in embryos lacking both the CE and DRR. (A-C) *MyoD* is expressed at E9.75 in the first branchial arch of wild-type embryos (A), but not in *MyoD^ΔCD/ΔCD^*(B) or *MyoD^ΔC18D/ΔC18D^* (C) embryos. Somite staining is evident in mutant embryos at this stage, but is reduced in intensity relative to wild-type embryos. **(D-F)** At E10.5, branchial arch staining and very faint forelimb bud staining is observed in mutant embryos (E, F). **(G-I)** At E11.5, the pattern of *MyoD* expression is very similar in wild-type and mutant embryos except that hindlimb bud staining is weaker in mutant embryos (H, I). **(J-L)** At E12.5, consistent differences in the pattern of *MyoD* expression between wild-type and mutant embryos is no longer discernible. In all panels, colored arrowheads represent expression, white arrowheads outlined in the same color represent lack of expression of the same structure: blue = mandibular arch; red = myotomes; black = limb buds; magenta = hypoglossal cord.

### Pax3-dependent MyoD expression occurs in the absence of the CE and DRR

In the trunk, *Myf5*, *Mrf4*, and *Pax3* function genetically upstream of *MyoD*, and in the absence of *Myf5* and *Mrf4*, *MyoD* expression in the trunk is dependent on *Pax3* (Kassar-Duchossoy et al., 2004; Tajbakhsh et al., 1997). We previously showed (Chen et al., 2002; Chen and Goldhamer, 2004) that embryos lacking either the CE or DRR remain responsive to *Pax3* when crossed to the original *Myf5*-null allele (Braun et al., 1992) (designated *Myf5^neo^* here), which likely also substantially reduced or eliminated expression of the closely linked *Mrf4* gene (see Kassar-Duchossoy et al., 2004). To test whether *Pax3*-dependent *MyoD* expression requires either the CE or the DRR, *MyoD^ΔCD/ΔCD^*;*Myf5^neo/neo^* embryos were generated, and *MyoD* expression was visualized by in situ hybridization. We utilized the original *Myf5* KO allele to maintain consistency with our previously published work (Chen et al., 2002; Chen and Goldhamer, 2004). As anticipated, overall levels of *MyoD* transcripts appeared reduced due to deletion of the CE and DRR (Fig. 3B, C). Importantly, however, *Pax3*-dependent recovery of *MyoD* expression in the trunk was observed, with expression phenocopying *MyoD* expression in *Myf5^neo/neo^* embryos (Fig. 3B, C, F, G). We also investigated whether sequences between the CE and DRR are required for *Pax3*-dependent rescue of *MyoD* expression. *MyoD^ΔC18D/ΔC18D^*;*Myf5^neo/neo^*embryos exhibited the same expression pattern as *Myf5^neo/neo^* and *MyoD^ΔCD/ΔCD^*;*Myf5^neo/neo^* embryos (Fig. 3D, H), demonstrating that the 18 kb sequence separating the CE and DRR is not required for *Pax3*-dependent expression of *MyoD*.

**Figure 3.**
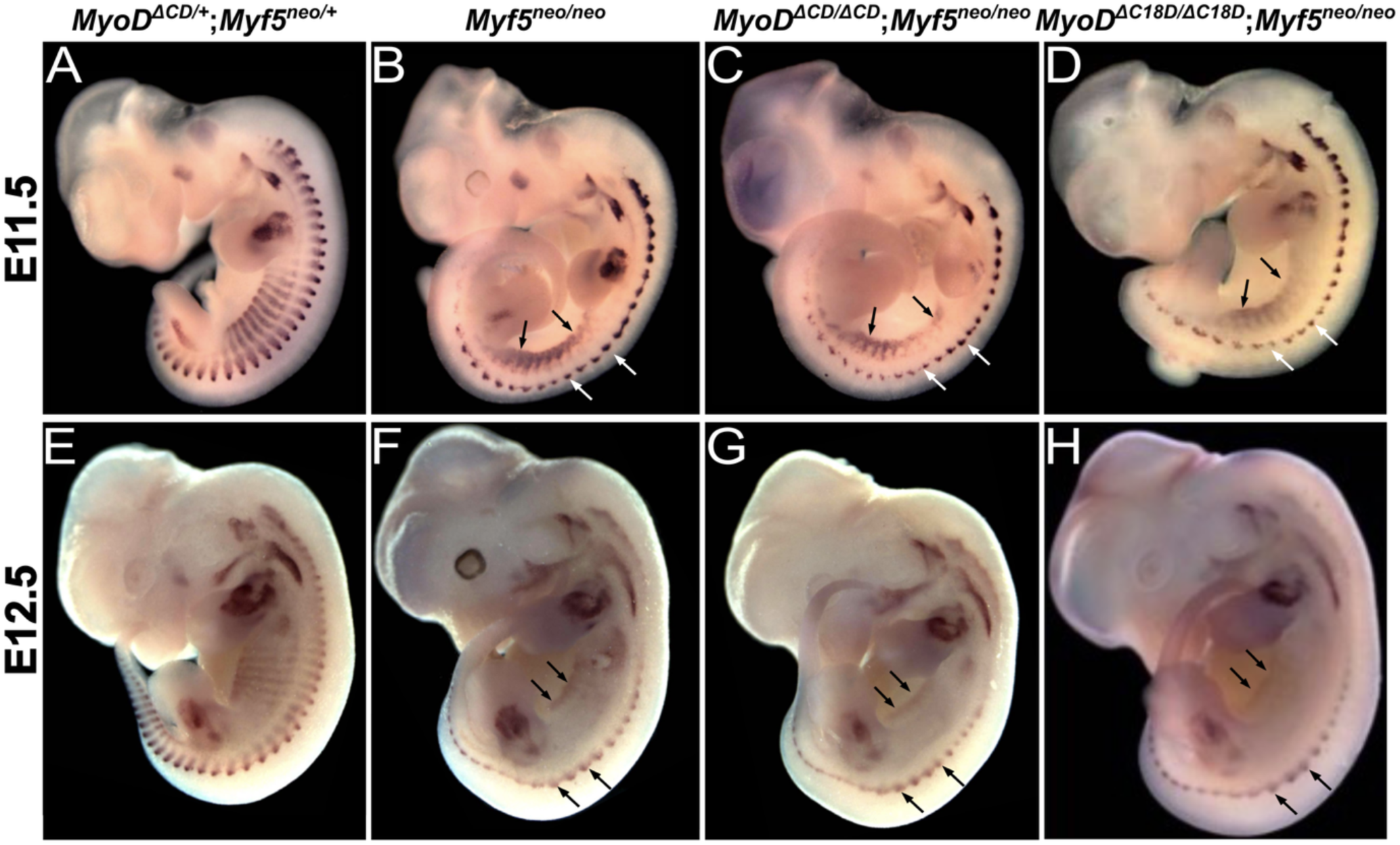
*Pax3*-dependent rescue of *MyoD* expression in *Myf5^neo/neo^* embryos does not require the CE, DRR, or sequences between them. (A, B, E, F) *MyoD* expression in the central myotomes is diminished in *Myf5^neo/neo^*embryos (which also likely eliminates *Mrf4* expression) at E11.5 (B) and E12.5 (F). Myotomal staining that remains at these stages is dependent on *Pax3*. **(C, D, G, H)** The staining pattern in the enhancer KO embryos is similar to that of *Myf5^neo/neo^* embryos (B, F). Arrows represent *Pax3*-dependent myotomal rescue.

### Computational analyses identify three candidate enhancer regions upstream of MyoD

The analyses above indicated that additional regulatory elements control *MyoD* expression during development. PRO-seq (Mahat et al., 2016) together with discriminative regulatory element detection from GRO-seq (dREG) (Danko et al., 2015) was used to identify putative transcription regulatory elements (TREs). dREG identifies regions of bidirectional transcription, which are associated with most active enhancers and promoters (Core et al., 2014; Danko et al., 2015). Cultured muscle stem cells (satellite cells) were used for this analysis because they are easily isolated in large numbers and high purity, and their growth and differentiation status can be easily controlled by manipulating culture conditions. As anticipated, the CE and DRR were identified as TREs by dREG. We note that the CE is contained within a larger dREG region that includes the enhancer region termed fragment 3 (Chen et al. 2001; Goldhamer et al. 1992; Goldhamer et al. 1995), which may contain regulatory information in addition to the CE (see Goldhamer et al. 1995; Chen et al. 2001). In addition to the CE and DRR, putative TREs were identified at approximately -36 kb and -96 kb relative to the *MyoD* TSS (Fig. 4; Supplemental Table 2). The putative TRE at -96 kb is embedded within the *Otog* gene, which regulates inner ear development (Schraders et al., 2012). As shown using publicly available chromatin accessibility and ChIP-seq datasets (Supplemental Table 3), these regions exhibit other features of regulatory regions, including DNase I hypersensitivity and enrichment for H3K27ac (Fig. 4). These regions also contain ChIP signals for the MRFs, a common feature of muscle-specific enhancers (Fig. 4). While not identified as a putative TRE by dREG, a region approximately 60 kb upstream of the *MyoD* TSS exhibits prominent ChIP signals for the MRFs, shows DNase I hypersensitivity, and is enriched for H3K27ac (Fig. 4). Survey of sequences up to approximately 250 kb upstream of the *MyoD* TSS did not identify additional regions with a strong PRO-seq signal or with ChIP signals comparable to the regions described above (data not shown). These results identify the -36, -60, and -96 kb regions as strong candidates for novel regulatory elements of *MyoD*.

**Figure 4.**
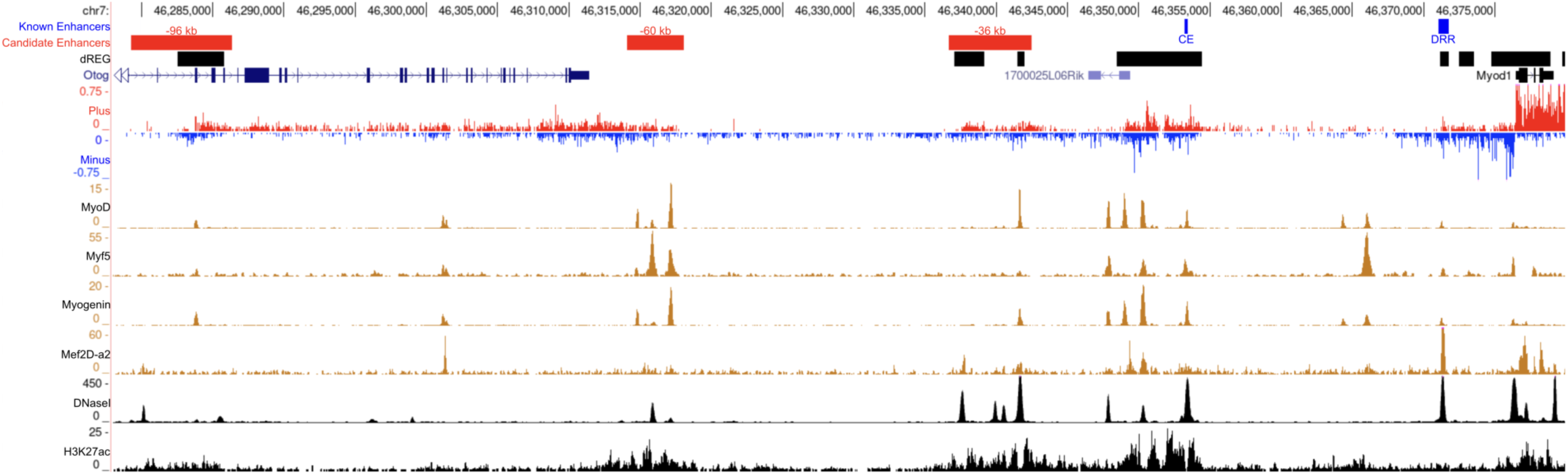
Computational analyses identify three candidate enhancer regions upstream of the *MyoD* core enhancer. Mouse *MyoD* and upstream sequence to approximately -100 kb are represented. PRO-seq analysis of cultured satellite cells was used to map polymerase density on the plus (red) and minus (blue) strands across the region. Known (CE and DRR) and candidate (-96 kb, -60 kb, and -36 kb) enhancer regions are represented by blue and red bars, respectively. Both the CE and DRR show bidirectional transcription (peaks on both the positive and negative strand), as identified by dREG (represented by the black bars on dREG track). dREG also identified the -96 kb and -36 kb regions as candidate enhancers. Areas of DNase I hypersensitivity and ChIP signals for H3K27ac and muscle regulatory factors are displayed. The -60 kb region was not identified by dREG, but was also chosen for functional testing based on its DNase I hypersensitivity and ChIP-seq profile. 1700025L06Rik refers to *LncMyoD*, and *Otog* is a regulator of inner ear development. Note the different scales for the ChIP-seq tracks.

### The -36 kb and -60 kb regions drive muscle-specific reporter gene expression in transgenic mice

The putative regulatory regions identified above were tested for their ability to drive muscle-specific *lacZ* expression in transgenic mouse embryos from E9.5 through E12.5. Two to three independent transgenic lines were created for each of the three regions to distinguish transcriptional activity attributable to each construct from site-of-integration effects. The relative length and position of the tested fragments and their relationship to the putative TREs at -96 kb and -36 kb are shown in Figure 4. The genomic coordinates and lengths of the tested fragments and dREG regions are provided in Supplemental Table 2. None of the three independently-derived -96lacZ lines exhibited muscle-specific expression at any stage tested; each line showed a distinct, non-muscle pattern of expression that is likely attributable to site-of-integration effects (data not shown). Although the putative TRE at -96 kb was not sufficient to drive *lacZ* expression, Hi-C data below is consistent with a role in *MyoD* regulation.

In contrast to -96lacZ transgenic embryos, -36lacZ and -60lacZ embryos exhibited expression patterns that together recapitulated most, but not all, aspects of *MyoD* expression. At E9.5-9.75, weak staining driven by -36lacZ was present in the anterior myotomes but absent from the ventral region of the interlimb somites, which typifies endogenous *MyoD* expression in E9.5 to E9.75 embryos (Fig. 5 A, B) (Chen et al. 2002; Chen and Goldhamer, 2004). Whereas *MyoD* is weakly expressed at E9.5 in the first branchial arch (Fig. 5A), -36lacZ expression was not detected at this stage (Fig. 5B). At E10.5, -36lacZ transgenic mouse embryos displayed a strong signal in the central myotomes along the anteroposterior embryonic axis, but expression was absent from the dorsal and ventral myotomal domains (Fig. 5D, E). Like endogenous *MyoD* mRNA, lacZ expression was observed in the forelimb buds at E10.5 (Fig. 5D, E). However, the staining pattern was broader and extended more distally than the signal for *MyoD* mRNA, indicating some ectopic staining in the forelimb buds (Fig. 5D, E). Weak staining was evident in the first branchial arch by E10.5, but lacZ expression was not detected in the cucullaris anlage and second branchial arch (Fig. 5D, E). At E11.5, -36lacZ was now expressed in most myogenic areas, including the cucullaris anlage and second branchial arch, indicating a delay in activation rather than a lack of domain-specific enhancer activity. Like at E10.5 (Fig. 5D, E), the anteroposterior extent of -36lacZ expression in the myotomes was similar to *MyoD* expression, although -36lacZ activity remained undetectable in the dorsal myotomal domain along much of the anteroposterior axis (Fig. G, H). At E12.5, essentially all myogenic regions expressed the -36lacZ transgene, except for the dorsal-most aspect of the epaxial myotomes (Fig. 5K).

**Figure 5.**
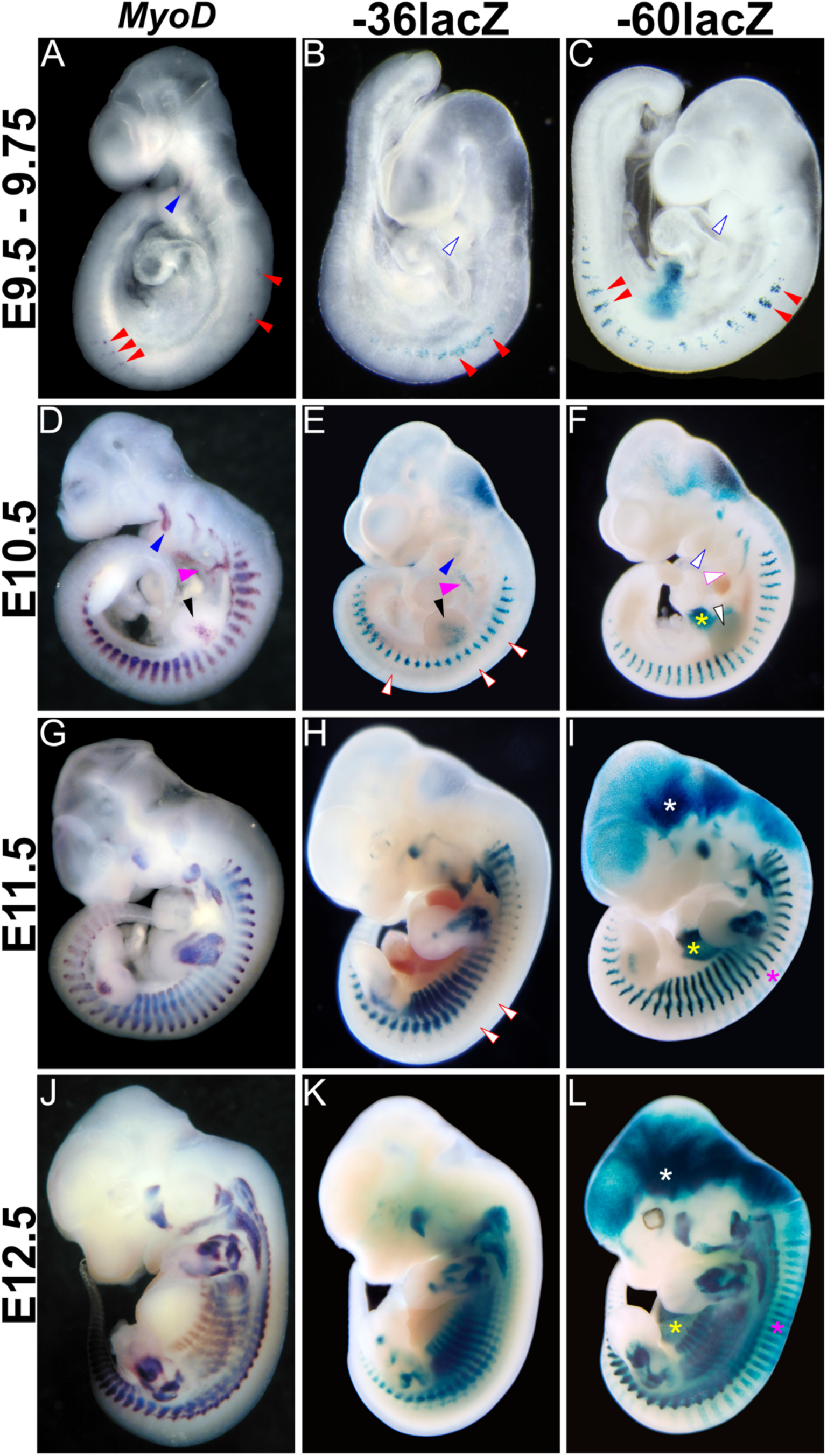
The -36 kb and -60 kb regions exhibit muscle-specific activity during development. Transgene expression was compared to embryos processed by whole-mount in situ hybridization for *MyoD* mRNA (first column). **(A-C)** Both the -36lacZ and -60lacZ transgenes show timely activation in somites. Note that the *MyoD* embryo (A) is younger than the transgenic embryos, accounting for the very weak staining. Staining in the mandibular arch is detectable in the *MyoD* embryo (A), but not the transgenic embryos. **(D-F)** At E10.5, *MyoD* is expressed in most myogenic centers, including the forelimb buds and hypoglossal cord. The -36lacZ transgene is weakly expressed in the mandibular arch and hypoglossal cord (E), but is not expressed in the dorsal aspects of the somites. Expression of the -60lacZ transgene (F) is not detectable in the mandibular arch, hypoglossal cord, or forelimb buds at this stage (F). **(G-L)** At E11.5 and E12.5, expression of both the -36lacZ and -60lacZ transgenes is similar to endogenous *MyoD* expression, except that expression of -36lacZ in dorsal somite domains remains absent. The - 60lacZ line exhibits ectopic expression in the head (white asterisks), dorsal trunk (vertical stripes in I and L; region denoted by pink asterisks), and liver (yellow asterisks); non-muscle expression was specific to this founder line and is likely due to site-of-integration effects. In all panels, colored arrowheads represent expression, white arrowheads outlined in the same color represent lack of expression of the same structure: blue = mandibular arch; red = myotomes; black = limb buds; magenta = hypoglossal cord.

The -60 kb region also exhibited muscle-specific activity in essentially all myogenic regions, but with a spatiotemporal pattern that deviated from both the -36 kb region and *MyoD*. At E9.5-9.75, staining was prominent in myotomes and included most interlimb somites (Fig. 5C). Like -36lacZ, expression was not observed in the branchial arches at this stage and was first observed at E10.5, although in both the - 36lacZ and -60lacZ transgenic lines, staining tended to be extremely weak and was undetectable in some embryos. Whereas -36lacZ transgenic embryos exhibited apparent muscle staining in the forelimb buds at E10.5, forelimb bud staining in -60lacZ embryos was delayed until E11.5 (Fig. 5E, F, I). Myotomal staining in -60lacZ embryos at E10.5 resembled *MyoD* expression; in contrast to -36lacZ embryos, staining was not restricted to the central myotome (Fig. 5D-F). At both E11.5 and E12.5, muscle staining of -60lacZ transgenic embryos closely resembled the expression pattern of *MyoD* (Fig. 5G, I, J, L). These results further support the notion that the -36 and -60 kb regions are novel *MyoD* enhancers.

### The -36 kb and -60 kb enhancer regions exhibit different dependencies on the MRFs

We next tested whether the -36 kb and -60 kb regulatory regions are responsive to auto-or cross-regulation by the MRFs. The -36lacZ and -60lacZ transgenes were crossed into MRF-deficient backgrounds (Table 1) and lacZ expression was assessed at E11.5. We used the *MyoD* null allele *MyoD^iCre^*, in which iCre replaces the entire coding sequence of exon 1 and a portion of intron 1 (Kanisicak et al., 2009). Two *Myf5* null alleles were used: *Myf5^loxP^*, which does not affect expression of the closely linked *Mrf4* gene (Kassar-Duchossoy et al. 2004), and *Myf5^neo^* (Braun et al. 1992), which likely eliminates expression of *Mrf4* (Table 1; see Kassar-Duchossoy et al. 2004). Expression of the -36lacZ and -60lacZ transgenes in a *Myf5^neo/+^*;*MyoD^iCre/+^*background was comparable to controls (compare Fig. 5H, I to Fig. 6A, C). -36lacZ and -60lacZ transgenic embryos lacking *MyoD* alone displayed a similar expression pattern to those on a WT background (Fig. 6A-D), indicating that *MyoD* is not essential for the activation of these enhancer regions or for maintenance of their activity. In embryos lacking *Myf5* alone, there was a modest reduction in lacZ staining in the central myotome, and the ventral extent of myotomal staining may have been slightly reduced (Fig. 6E, G).

**Figure 6.**
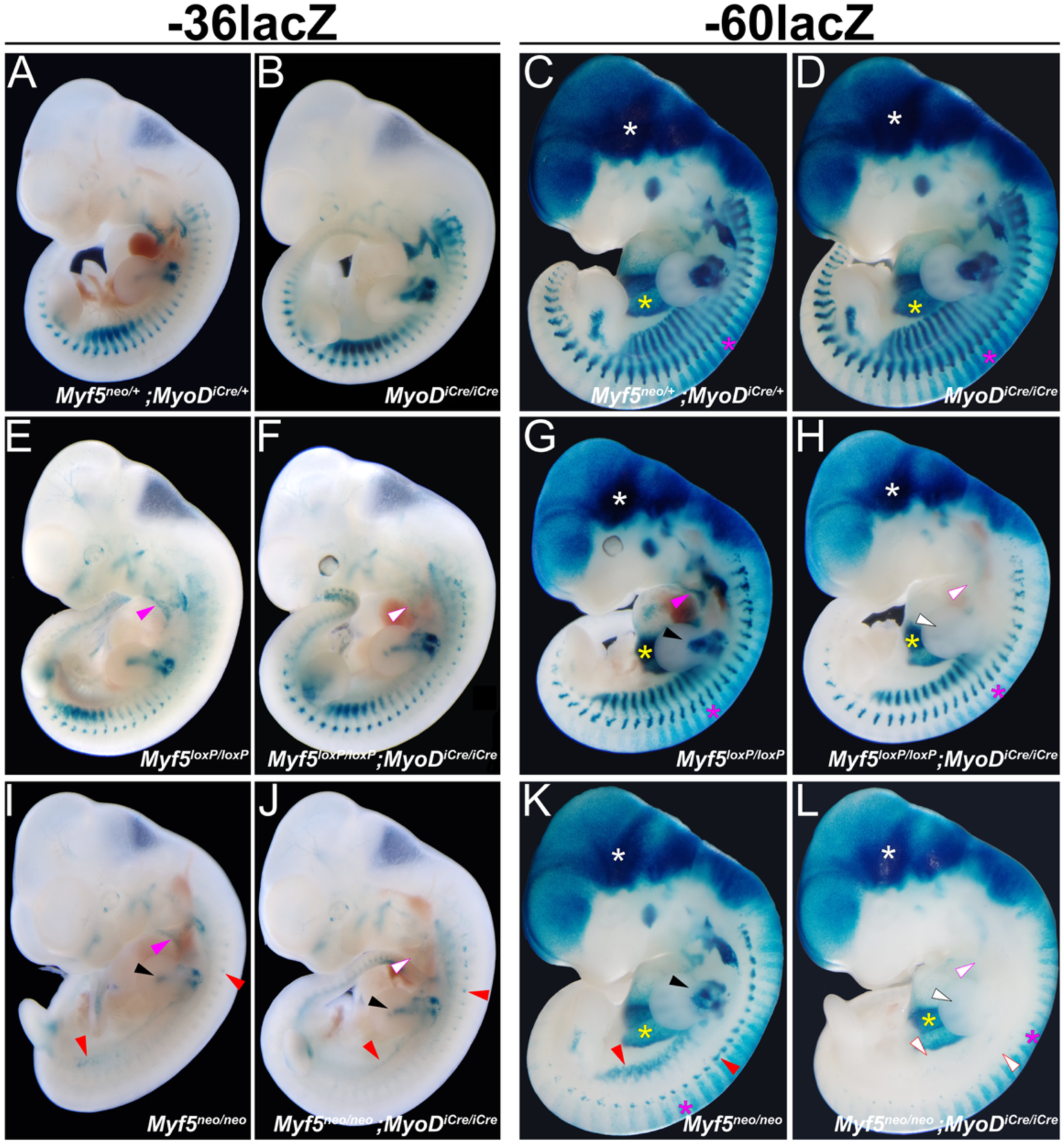
The -36 and -60 enhancer regions exhibit differential dependencies on the MRFs. Transgene expression was compared at E11.5. **(A-D)** Expression of -36lacZ (B) and -60lacZ (D) are not appreciably affected in a *MyoD* null (*MyoD^iCre^*^/*iCre*^) background. **(E, F)** In a *Myf5* null (*Myf5^loxP^*^/*loxP*^) background, the pattern of -36lacZ expression (E) is similar to controls (A), although staining intensity may be slightly weaker, particularly in central myotomes. In -36lacZ embryos lacking both *Myf5* and *MyoD* (F), expression in the hypoglossal cord is diminished or absent, while other aspects of transgene expression resemble *Myf5^loxP^*^/*loxP*^ embryos. **(G, H)** As with -36lacZ, consistent changes in -60lacZ expression are not obvious in *Myf5^loxP^*^/*loxP*^ embryos, except for a slight reduction in staining in central myotomes. Unlike -36lacZ, in addition to loss of hypoglossal cord staining, limb bud staining is absent in embryos lacking both *Myf5* and *MyoD* (H). **(I, J)** Myotomal expression of -36lacZ is substantially reduced, but not eliminated, in *Myf5^neo/neo^* embryos (I), and the loss of *MyoD* (J) does not reduce transgene expression further. **(K, L)** Like - 36lacZ, expression of -60lacZ is reduced in the myotomes in *Myf5^neo/neo^*embryos (K). The additional loss of *MyoD* abrogates myotomal and limb bud expression (L). Faint staining in the head and neck persisted. Ectopic expression in -60lacZ embryos is not affected by MRF status. In all panels, colored arrowheads represent expression, white arrowheads outlined in the same color represent lack of expression of the same structure: red = myotomes; black = limb buds; magenta = hypoglossal cord. In this -60lacZ line, there is ectopic expression in the head (white asterisk), dorsal trunk (vertical stripes), and liver (yellow asterisk).

**Table 1.**
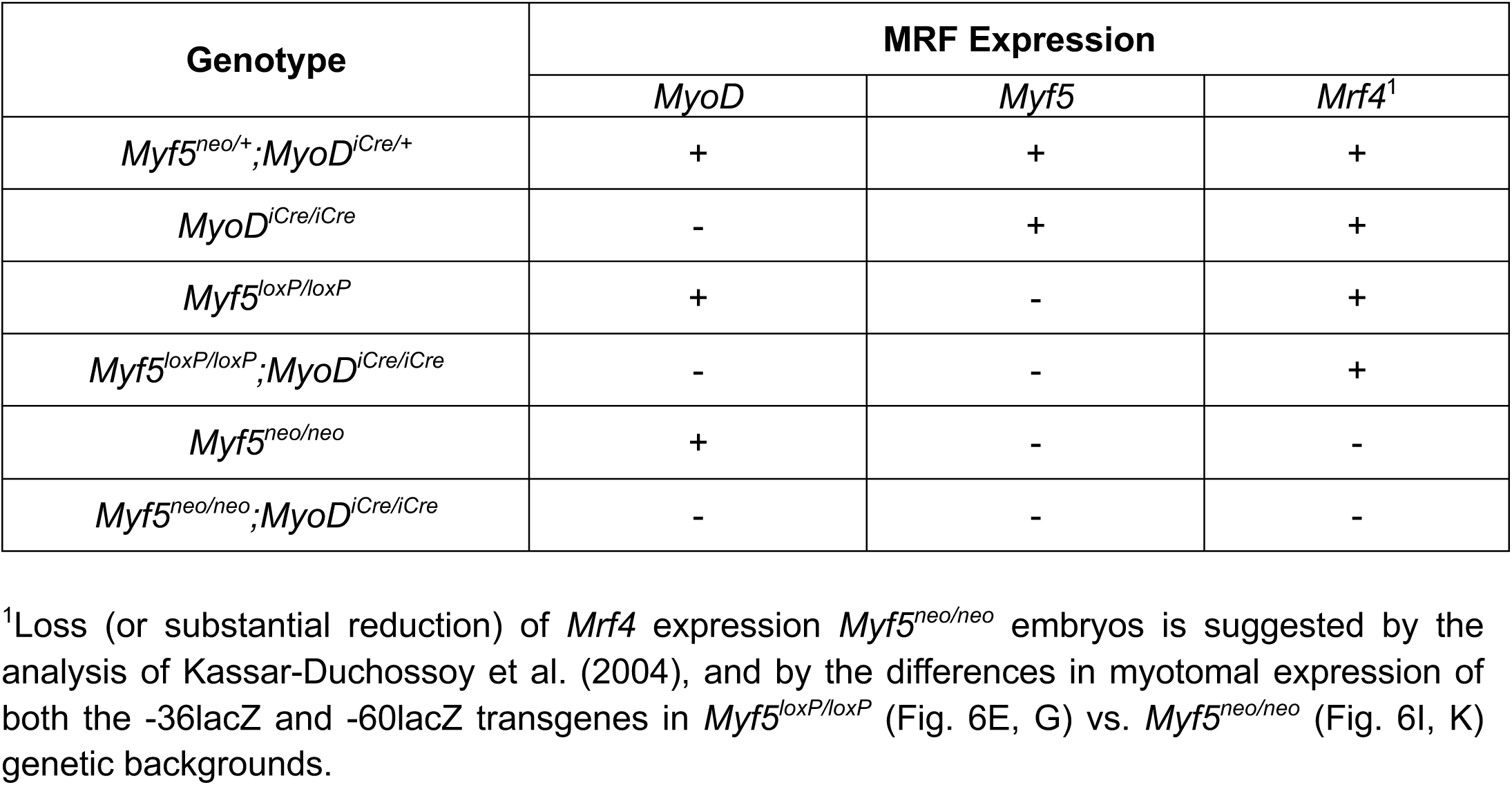
MRF Expression in different MRF-deficient backgrounds.

| Genotype | MRF Expression |  |  |
| --- | --- | --- | --- |
|  | *MyoD* | *Myf5* | *Mrf4*<sup>1</sup> |
| *Myf5*<sup>neo/+</sup>;*MyoD*<sup>iCre/+</sup> | + | + | + |
| *MyoD*<sup>iCre/iCre</sup> | - | + | + |
| *Myf5*<sup>loxP/loxP</sup> | + | - | + |
| *Myf5*<sup>loxP/loxP</sup>;*MyoD*<sup>iCre/iCre</sup> | - | - | + |
| *Myf5*<sup>neo/neo</sup> | + | - | - |
| *Myf5*<sup>neo/neo</sup>;*MyoD*<sup>iCre/iCre</sup> | - | - | - |

<sup>1</sup>Loss (or substantial reduction) of *Mrf4* expression *Myf5*<sup>neo/neo</sup> embryos is suggested by the analysis of Kassam-Duchossoy et al. (2004), and by the differences in myotomal expression of both the -36lacZ and -60lacZ transgenes in *Myf5*<sup>loxP/loxP</sup> (Fig. 6E, G) vs. *Myf5*<sup>neo/neo</sup> (Fig. 6I, K) genetic backgrounds.

Differential effects of MRF deficiencies on the activities of the -36 kb and -60 kb regions became apparent in embryos lacking both *MyoD* and *Myf5* (*Myf5^loxP/loxP^*;*MyoD^iCre/iCre^*). In addition to the minor deficits observed in a *Myf5^loxP/loxP^* background, -36lacZ was not expressed in the hypoglossal cord, which contributes to musculature of the tongue (Fig. 6F). -60lacZ embryos in this genetic background also lacked expression in the hypoglossal cord (Fig. 6H). Further, limb bud expression was absent at E11.5 (Fig. 6H), indicating that either *Myf5* or *MyoD* is required for the timely activation of the -60 kb region in both the forelimb and hindlimb buds. The deficit was temporary, however, as -60lacZ expression in limb buds was observed at E13.5 (data not shown). -36lacZ and -60lacZ expression was reduced further in embryos deficient for both *Myf5* and *Mrf4* (*Myf5^neo/neo^* embryos), with the myotomes being most affected (Fig. 6E, G, I, K). Expression of -36lacZ in the myotomes of *Myf5^neo/neo^* embryos was almost completely eliminated, while staining in the branchial arches approximated that of *Myf5^loxP/loxP^*embryos. In a *Myf5^neo/neo^* background, expression of -60lacZ persisted in the ventral myotomal domain of interlimb somites and the dorsal myotomal domain of all somites. Because the overall staining intensity of the - 60lacZ line was greater than the -36lacZ line, it is unclear whether relative differences between these lines are biologically meaningful. In both transgenic lines, staining persisted, albeit at slightly lower intensities compared to *Myf5^loxP/loxP^* embryos, in muscle regions derived from migratory lineages, including the limb buds, branchial arches, and hypoglossal cord (Fig. 6E, G, I, K). Finally, transgene expression was assessed in E11.5 embryos deficient in all three myogenic factors (*Myf5^neo^*^/*neo*^;*MyoD^iCre^*^/*iCre*^ embryos) to test whether transgene activity that persisted in *Myf5^neo^*^/*neo*^ embryos was dependent on *MyoD*. Expression of the -36lacZ transgene on this genetic background closely resembled its activity in *Myf5^neo^*^/*neo*^ embryos (Fig. 6I, J). In contrast, the additional loss of *MyoD* essentially abrogated all remaining activity of the -60 kb element, except for very weak expression in the branchial arches (Fig. 6L). Collectively, genetic analyses indicate that activity of the -36 kb region does not require *MyoD*, and its activity in limb buds and branchial arches is largely independent of all three MRFs tested. In contrast, activity of the -60 kb element in limb buds requires either *Myf5* or *MyoD*. Finally, myotomal activity of both the -36 kb (Fig. 6E, I) and -60 kb (Fig. 6G, K) elements is highly dependent on either *Myf5* or *Mrf4*, and the additional loss of *MyoD* abrogates the remaining myotomal activity of the -60 kb region (Fig. 6K, L).

### The -36 kb enhancer is active in adult satellite cells

To begin to test whether the newly identified *MyoD* regulatory elements are active at adult stages, we tested whether the -36lacZ and -60lacZ transgenes are active in cultured satellite cells isolated from muscle of adult mice. Satellite cells were isolated from the hindlimb muscles of -36lacZ and -60lacZ transgenic mice using MACS and assayed for lacZ expression under both growth and differentiation conditions. Satellite cells harboring the -24lacZ transgene, which contains 24 kb of human *MyoD* 5’ flanking sequence, including the CE and DRR (Goldhamer et al. 1992; Chen et al. 2001), served as a comparison group. Both the -24lacZ and -36lacZ transgenes were expressed in satellite cell cultures (Supplemental Fig. 2). In contrast, only rare lacZ-positive satellite cells were observed in cultures derived from -60lacZ mice (Supplemental Fig. 2). The lack of apparent activity of the TRE at -60 kb in cultured satellite cells may explain why this region was not identified by dREG enhancer prediction software.

### 3D chromatin architecture of the MyoD locus: cis-regulatory interactions and topologically associating domains

Evidence presented above strongly implicates novel regulatory regions in the control of *MyoD* expression. We next explored the chromatin architecture around these regions and the *MyoD* gene to establish whether these regions interact in *cis* with each other or *MyoD*. Indeed, analysis of Hi-C data from primary myoblasts cultured under both growth and differentiation conditions (Wang et al., 2022) revealed a topologically associating domain (TAD) encompassing the genomic region from the -96 kb region to *MyoD*. All three candidate enhancers are involved in multiple interactions with each other, the CE, and *MyoD* under both growth and differentiation conditions, and with the DRR under differentiation conditions (Fig. 7). The data also suggest interactions between the CE and DRR under differentiation conditions. These data are consistent with the PRO-seq and transgenic data and further suggest that an enhancer hub may form with multiple interactions throughout this domain (Fig. 7). Under both growth and differentiation conditions, only the outermost loop, involving the -96 kb region and *MyoD*, is associated with CTCF at both anchors, while the remaining loops inside this domain are mediated by *cis*-regulatory interactions (Fig. 7). Notably, this highly organized chromatin architecture is absent in mESCs, which do not express *MyoD*, demonstrating that these looping interactions are not a general feature of this chromosomal locus (Supplemental Fig. 3). Whether these interactions are restricted to myogenic cells will require further investigation.

**Figure 7.**
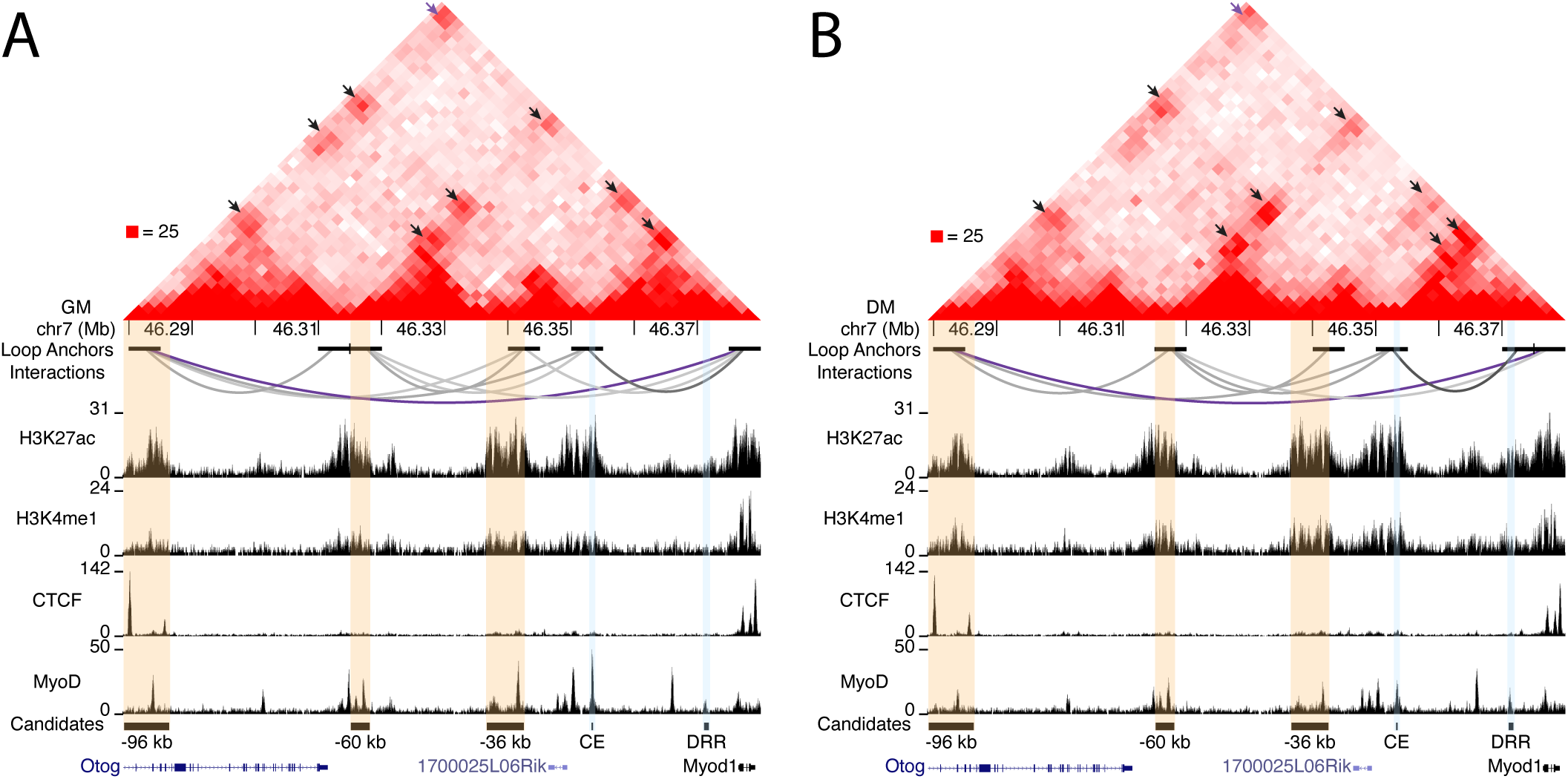
Cis-regulatory interactions within a topologically associating domain that encompasses candidate regulatory elements, known enhancers, and *MyoD*. (Top) Hi-C contact map of the *MyoD* locus in myoblasts cultured in growth (A) and differentiation (B) media. Interaction peaks are indicated by arrows and maximum intensity is shown on the left. (Middle) Individual loop anchors and anchor interactions are shown as arcs. (Bottom) ChIP-seq profiles for H3K27ac, H3K4me1, CTCF, and *MyoD*, and coordinates of validated candidate (orange) and known (light blue) enhancers and genes. These enhancers overlap with histone modifications associated with active enhancers and loop anchors that indicate contact with *MyoD*. The CTCF-delimited loop between the -96 kb region and *MyoD* is shown in purple.

Taking a broader view of the *MyoD* locus, we identified another region of interest 220 kb downstream of *MyoD*, within intron 8 of the *Sergef* gene, which is widely expressed. Overlapping H3K27ac, H3K4me1, and CTCF ChIP-seq signals coincide with a loop anchor that interacts with the -96 kb region under both growth and differentiation conditions, as well as with *MyoD* and the -60 kb element under growth conditions (Supplemental Fig. 4). In addition, multiple contact points residing between *MyoD* and the +220 kb region were identified. Functional analyses are required to determine whether these regions represent bona fide *MyoD* regulatory elements.

## DISCUSSION

Myoblast identity in the embryo is established through the coordinated action of *MyoD*, *Myf5* and *Mrf4*, each of which is expressed in a distinct spatiotemporal pattern during development. Detailed transgenic studies of *Myf5* and *Mrf4 cis* regulation have demonstrated a high degree of regulatory modularity and synergy such that the collective activity of many enhancers is required to recapitulate the dynamic expression pattern of these genes in the embryo (Carvajal et al., 2001; Hadchouel et al., 2000; Summerbell et al., 2000; Teboul et al., 2002; Zweigerdt et al., 1997). This complexity is perhaps not surprising given the diverse embryonic signaling environments within which muscle determining genes are activated (Hernández-Hernández et al., 2017; Lima and Relaix, 2021). By contrast, the regulation of *MyoD* was considered comparatively simple, as only two *MyoD* regulatory regions had been identified prior to the current study (CE and DRR) (Asakura et al., 1995; Goldhamer et al., 1995, 1992; Tapscott et al., 1992), and their combined activities could largely explain the timing and spatial patterning of *MyoD* expression in the embryo. However, the phenotype of embryos lacking both the CE and DRR demonstrated greater regulatory complexity than previously known, and we identified novel DNA elements that collaborate to regulate *MyoD* expression.

The CE and DRR have been extensively studied using both transgenic and knockout approaches, and important insights have come from comparisons of these approaches. These studies showed that the CE is both necessary (Chen and Goldhamer, 2004) and sufficient (Chen et al., 2001; Goldhamer et al., 1995; Kablar et al., 1998; Kucharczuk et al., 1999) for the timely activation of *MyoD* in migratory myogenic lineages of the branchial arches and limb buds. These data are consistent with DRR-lacZ transgene expression, which is delayed in these myogenic populations (Asakura et al., 1995; Chen et al., 2001; Kablar et al., 1998). The CE requirement for the timely activation of *MyoD* may be at least partly attributable to the enhancer RNA, ^CE^RNA (Mousavi et al., 2013), as the histone methyltransferase LSD1 is required for transcription of the ^CE^RNA, and conditional inactivation of *Lsd1* in mouse embryos disrupted the timely activation of *MyoD* in limb buds (Scionti et al., 2017). In contrast to its essential role in migratory muscle lineages, deletion of the CE did not affect the timing or pattern of myotomal expression of endogenous *MyoD* (Chen and Goldhamer, 2004), despite the identification of CE elements that are specifically required for enhancer activity in the myotomes of transgenic mice (Kucharczuk et al., 1999). Further, while *MyoD* expression in the trunk in *Myf5^nlacZ^* KO mice (which also eliminates *Mrf4* expression; see Kassar-Duchossoy et al. 2004) requires *Pax3* (Tajbakhsh et al., 1997), single enhancer KO studies showed that both the CE (Chen and Goldhamer, 2004) and DRR (Chen et al., 2002) are dispensable for *Pax3*-dependent compensatory activity. These data are consistent with transgenic studies showing that both the CE and DRR exhibit activity in *Myf5*-mutant embryos (Kablar et al., 1999, 1998, 1997). Collectively, these data raised the non-mutually exclusive possibilities that the CE and DRR exhibit partial functional redundancy, or that *MyoD* is also regulated by unknown enhancer elements.

We tested for functional redundancy between the CE and DRR by analyzing *MyoD* expression in KO mice lacking both enhancers. Interestingly, *MyoD* transcript levels were significantly reduced in dKO embryos compared to WT embryos (this study) and those lacking the CE or DRR alone (Chen et al., 2002; Chen and Goldhamer, 2004). While the CE and DRR lack extended regions of sequence similarity (Chen et al., 2001; Goldhamer et al., 1995; Kucharczuk et al., 1999), both elements contain multiple E-boxes (binding sites for the MRFs), and the significant reduction in *MyoD* expression levels may reflect disruption of auto-and cross-regulatory interactions with MRF family members required to maintain appropriate *MyoD* transcriptional levels. Future work is required to determine the extent to which the significant reduction in *MyoD* transcription is mediated by the loss of possible synergistic interactions between the ^CE^RNA and ^DRR^RNA/MUNC, both of which have been shown to regulate *MyoD* transcription (Mousavi et al., 2013; Mueller et al., 2015). Interestingly, PRO-seq data of cultured satellite cells lacking both *MyoD* and *Myf5* demonstrated reduced transcription from both the plus and minus strands in the DNA region encompassing the CE (unpublished observations), indicating that ^CE^RNA transcription is directly or indirectly regulated by one or both MRFs.

Despite synergy between the CE and DRR in regulating *MyoD* levels, spatiotemporal patterns of *MyoD* expression in dKO mice reflected the additive effects of the individual enhancer KOs. Further, expression deficits were not worsened when the ∼18 kb of sequence separating the CE and DRR was also deleted, indicating no evident regulatory role for these intervening sequences. These data indicate that functional redundancy between the CE and DRR cannot explain the persistent muscle specificity of *MyoD* expression exhibited in the single enhancer KOs. Additionally, compensatory mechanisms that regulate *MyoD* expression in *Myf5^neo/neo^* KO embryos were largely operative in dKO embryos. Collectively, these data strongly indicate that heretofore unknown DNA elements contribute to the *cis* regulatory landscape of *MyoD*.

We used PRO-seq and available ChIP-seq and DNase I datasets to identify putative regulatory regions of *MyoD*. We focused on three upstream regions, one of which (-60 kb) was not identified as a candidate enhancer using dREG, which identifies regions of bidirectional transcription, a characteristic of most active enhancers and promoters (Core et al., 2014; Danko et al., 2015). However, the -60 kb region is in an open chromatin configuration and includes prominent ChIP peaks for the MRFs, H3K4me1, and H3K27ac. At least one study has shown that transcription of eRNAs is a critical component of enhancer activity (Tippens et al., 2020); thus, the absence of both a detected TRE and enhancer reporter activity in cultured satellite cells suggests that activity of the -60 kb enhancer may be restricted to myogenic cells in the embryo. The -96 kb and -36 kb regions share these chromatin features with the -60 kb region and, additionally, were identified by dREG as potential TREs. Arguing against a role for the -96 kb region in regulating *MyoD* is the finding that none of the three independently-derived -96lacZ mouse lines showed muscle-specific expression in embryos. However, studies have identified DNA elements within super-enhancers that amplify transcriptional output but have no intrinsic activity when tested in isolation (Hay et al., 2016; Sahu et al., 2022). These elements have been termed facilitators (Blayney et al., 2023), and although their mechanism of action is not entirely clear, they may tether enhancers together, thereby facilitating their interaction (Levo et al., 2022). Notably, Hi-C analysis of chromatin architecture revealed that the -96 kb region interacts with other enhancers upstream of *MyoD*, consistent with the testable model that the -96 kb region functions as a facilitator of *MyoD* transcription.

The -36 kb and -60 kb DNA elements drove lacZ expression in the myotomes, limb buds, and branchial arches. Whereas their combined activities loosely approximated the spatiotemporal pattern of *MyoD* expression, their patterns of activity were distinct from each other and from the CE and DRR. These activity differences suggest that the -36 kb and -60 kb regions probably do not represent fully redundant shadow enhancers, which are common components of regulatory networks that control expression of developmental control genes in *Drosophila* (Cannavò et al., 2016). This is supported by the findings that CE/DRR dKO embryos exhibited substantially reduced *MyoD* levels, and that CE KO (Chen and Goldhamer, 2004) and dKO embryos showed deficits in *MyoD* initiation in migratory muscle lineages. We emphasize, however, that the -60 kb and -36 kb elements share similarities with the DRR and CE, respectively, and partial redundancy cannot be ruled out. Like the DRR (Kablar et al., 1999), the -60 kb element exhibited delayed activity in the limb buds and essentially complete dependency on MRF function in transgenic assays. CE and -36 kb elements are separately sufficient to drive the timely expression of lacZ in the limb buds, and both the CE (Kablar et al., 1999) and -36 kb enhancers retain partial activity in mice lacking all MRF function, consistent with roles in initiating *MyoD* expression. Systematic deletion analyses and functional studies of transcripts associated with the -60 kb and -36 kb regions are required to more clearly define the regulatory relationships between these four DNA elements.

The skeletal muscle-specific activity of the -60 kb and -36 kb DNA elements, their proximity to *MyoD*, and the absence of other known muscle genes in this genomic region provided strong evidence, but not proof, that these newly identified elements contribute to the transcriptional regulation of *MyoD*. Important supporting evidence came from analysis of high-resolution 3D chromatin architecture data from primary cultures of proliferating and differentiating myoblasts (Wang et al., 2022), analyses that showed combinatorial interactions between the upstream regulatory regions and with *MyoD*. mESCs, which served as a non-myogenic comparison, showed no such interactions in this genomic region. For both growth and differentiation conditions, Hi-C data reflect the total of interactions across the myogenic cell population at one point in time, and the extent to which the interactions observed reflect developmental heterogeneity within the population or a “snapshot” of a highly dynamic process shared by most cells is not clear. Interestingly, enhancer interactions detected in proliferating and differentiating cultures were distinct but overlapping. One noteworthy difference was that interactions between the CE and DRR were restricted to differentiating cells, consistent with known activities of the DRR in embryos. These data point to a high degree of regulatory complexity, even in this comparatively simple cell culture model of myogenesis.

*MyoD* was among the first genes identified that regulate lineage determination in vertebrates, and it serves as an important model to understand how upstream signaling events in the embryo are integrated to control the expression of genes that determine cell identity. The present study strongly argues that at least four DNA elements regulate the developmental expression of *MyoD*. It is likely, however, that additional elements and interactions are involved in *MyoD* cis-regulatory control. While we focused on enhancers upstream of *MyoD*, Hi-C also detected multiple looping interactions involving sites 3’ of *MyoD*, and the number of such contact points varied between growth and differentiation conditions. Notably, all identified interacting regions upstream and downstream of *MyoD* were contained within a CTCF-delimited loop spanning approximately 300 kb, from the -96 kb region to a contact point approximately 220 kb downstream of *MyoD*. Additional work is needed to fully understand how these known and putative enhancer regions interact to control the initiation and maintenance of *MyoD* expression. Recently, it was shown that MYOD serves as an anchor protein to organize the 3D genome organization of muscle cells, which may determine their myogenic identity (Wang et al., 2022). As *MyoD* is auto-and cross-regulated by the MRFs, it will be interesting to determine whether the 3D chromatin architecture of the *MyoD* genomic locus is itself regulated by MYOD and other MRFs.

## ACKNOWLEDGMENTS

We thank members of the Goldhamer lab, past and present, for their valuable input during the course of these studies. We also thank Dr. Adam Zweifach for advice on statistical analysis, UCONN Health’s Center for Mouse Genome Modification for generation of chimeric mice and pronuclear injections, the Flow Cytometry Facility for assistance with satellite cell isolation, and the Center for Genome Innovation for sequencing. This work was supported by funding from the University of Connecticut to J.E., and L.J.C., by grants from the NIH to J.E. (NIGMS; R35GM146922), L.J.C. (NIGMS; R35GM128857), and D.J.G. (NIAMS; R01AR44878 and R01AR076394), and by a grant from the Muscular Dystrophy Association to D.J.G.

