## Supplemental Materials for "Developmental expression of the skeletal muscle determination gene, *MyoD*, is regulated by novel enhancer elements that interact with the core enhancer and distal regulatory region"

### Supplemental Figures 1-4

Fig. S1: MyoD quantification

Fig. S2: Expression of transgenes in satellite cell culture

Fig. S3: Hi-C analysis of *MyoD* locus in mESCs

Fig. S4: Identification of potential enhancer elements downstream of *MyoD*

### Supplemental Tables 1-3

Table S1: Primers used for genotyping

Table S2: Coordinates of dREG candidate enhancers and regions cloned for transgenic analyses

Table S3: Hi-C and ChIP-seq datasets

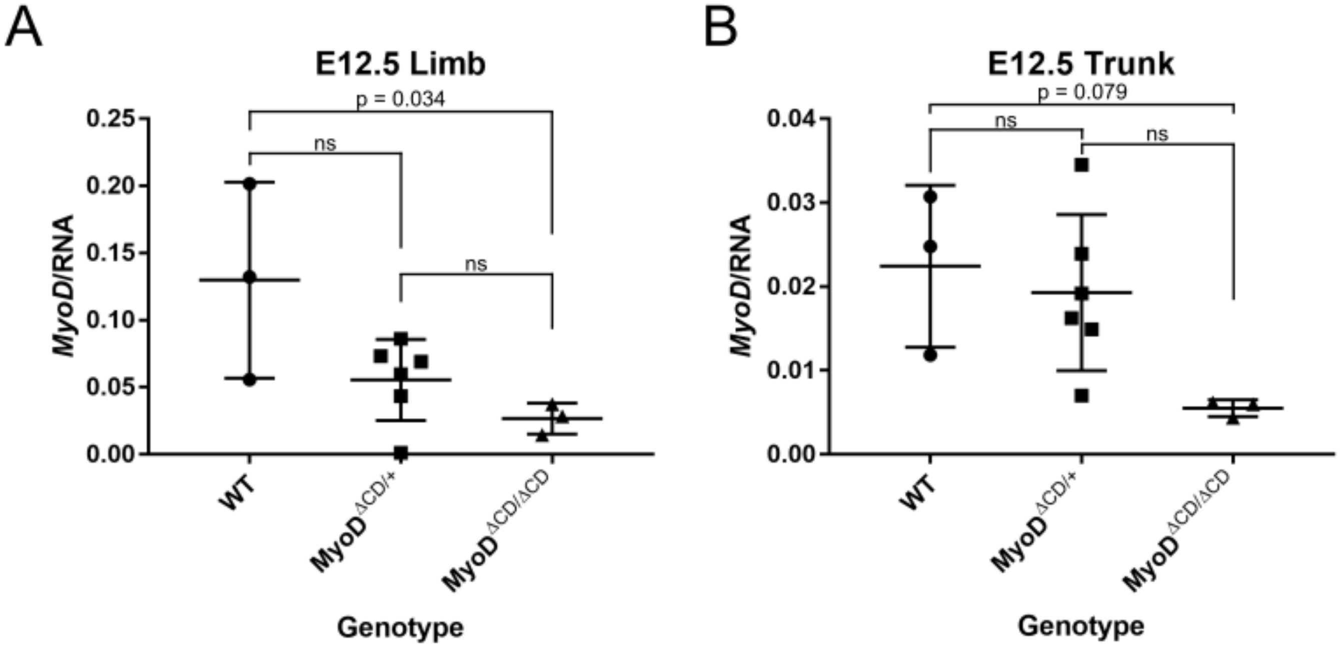


**Supplemental Figure 1. ddPCR comparing *MyoD* RNA between WT, *MyoD*$^{\Delta CD/+}$, and *MyoD*$^{\Delta CD/\Delta CD}$ embryos.** E12.5 embryos were harvested and either the limb buds or trunk was isolated for ddPCR. *MyoD* was normalized to input RNA. Significance was assessed by one-way ANOVA with Tukey's multiple comparisons.

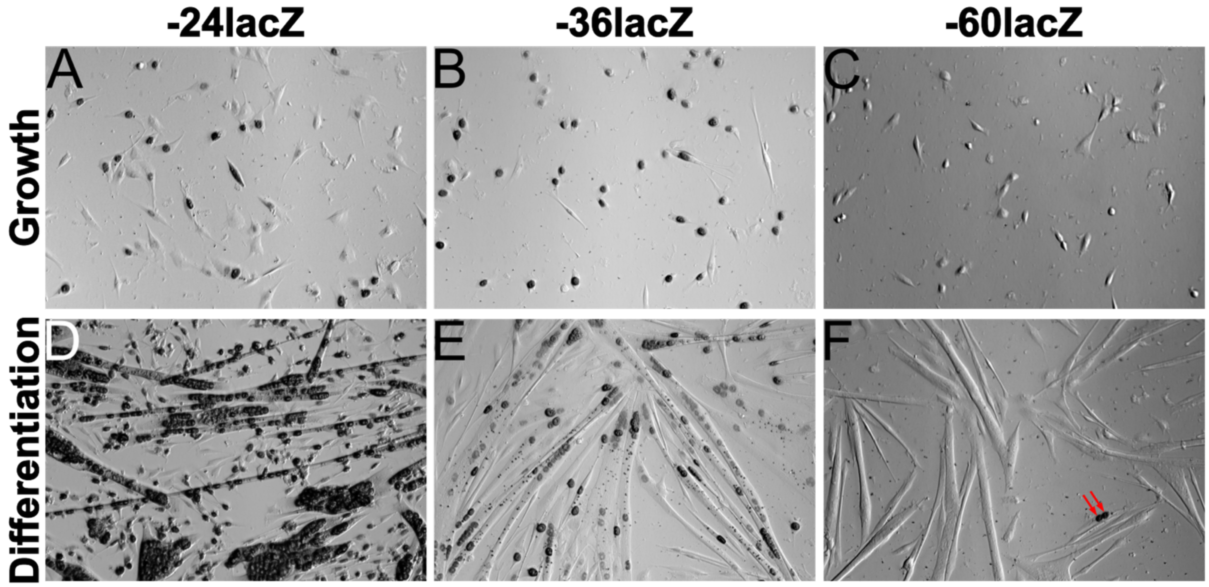


**Supplemental Figure 2. Expression of lacZ transgenes in satellite cell cultures.** Satellite cells from -24lacZ (contains both the CE and DRR), -36lacZ, and -60lacZ transgenic mice were isolated via MACS and cultured in either growth or differentiation media. lacZ-positive cells are numerous in -24lacZ and -36lacZ cultures under growth conditions (A, B). Essentially all myotubes derived from -24lacZ and -36lacZ satellite cells contain lacZ-positive nuclei under differentiation conditions (D, E). Whether X-gal staining in these myotubes represents transgene expression or the perdurance of β-gal is not known. X-gal-positive cells are rare under both growth and differentiation conditions in -60lacZ satellite cell cultures (C, F) (arrows show X-gal-positive nuclei in F). Note that the lacZ expression constructs contain a nuclear localization signal.

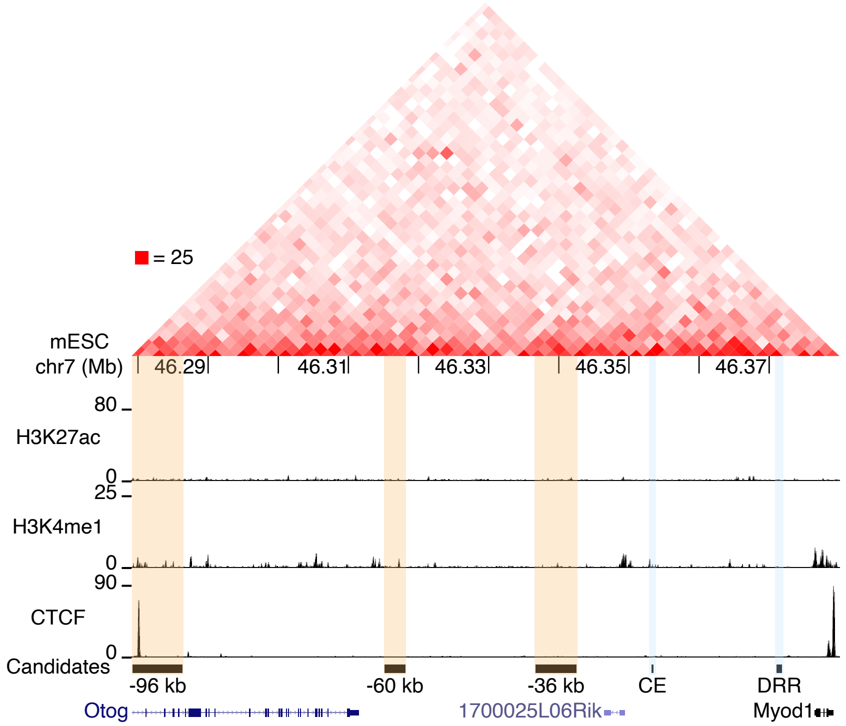


**Supplemental Figure 3. 3D genomic landscape of the *MyoD* locus and candidate enhancers in mESCs.** (Top) Hi-C contact map with maximum intensity shown on the left. (Middle) ChIP-seq profiles for H3K27ac, H3K4me1, and CTCF. (Bottom) Coordinates of validated candidate (orange) and known (light blue) enhancers, and genes. No looping interactions between enhancer regions or with *MyoD* were identified.

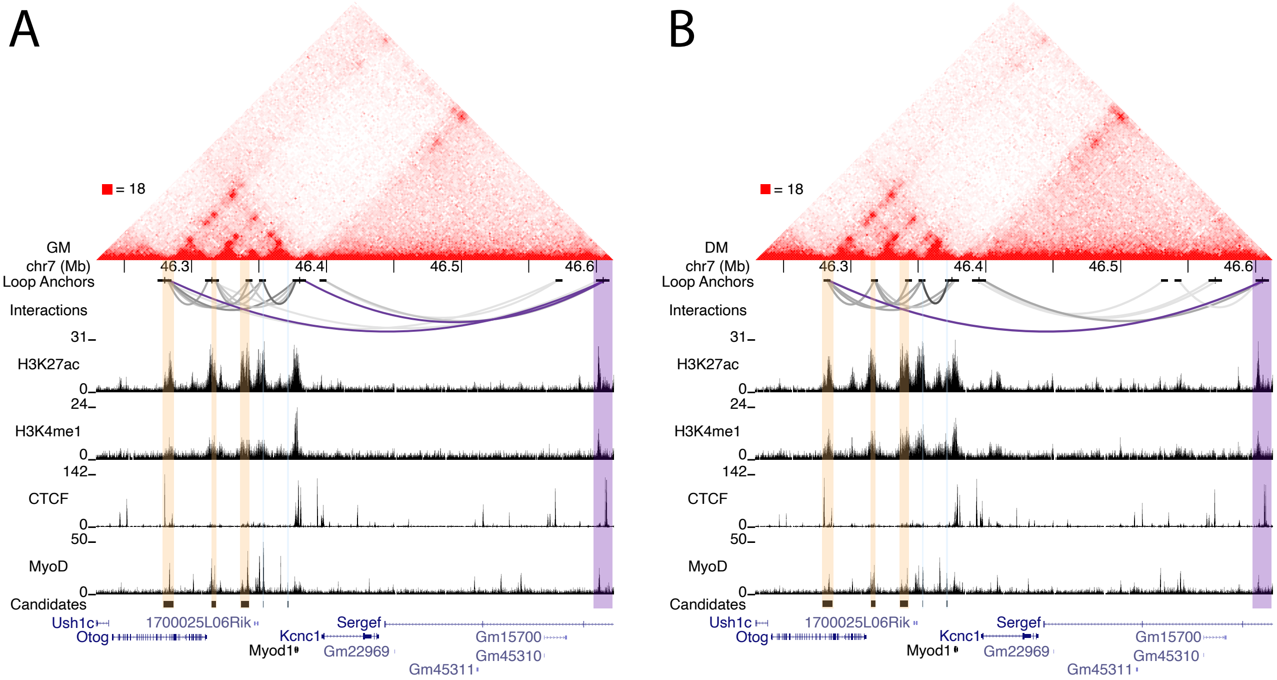


**Supplemental Figure 4. Identification of potential uncharacterized *MyoD* enhancer regions downstream of *MyoD*.** (Top) Hi-C contact map of the *MyoD* locus in myoblasts cultured in either growth (A) or differentiation (B) media. Maximum intensity is shown on the left. (Middle) Individual loop anchors and anchor interactions are shown as arcs. (Bottom) ChIP-seq profiles for H3K27ac, H3K4me1, CTCF, and *MyoD*. Coordinates of the -96 kb, -60 kb and -36 kb regions are shown in orange and coordinates of the CE and DRR are shown in light blue. Known genes in this genomic region are shown. Loops anchored by the regions at -96 kb and +220 kb (A, B) or between the +220 kb region and *MyoD* (A), are shown in purple.

**Supplemental Table 1. Primers used for genotyping**

| Allele | Forward/Reverse Primer | Size (bp) | PMID |
|---|---|---|---|
| *MyoD*$^{\Delta CD}$ | Forward 5'-CTTGGAACCACACTACCTCAAGG-3'<br>Reverse 5’-GTTCCTCTCATGCCTGGTGTTTAGG-3' | 230 | Present Study |
| *MyoD*$^{\Delta C18D}$ | Forward 5'-CTTGGAACCACACTACCTCAAGG-3'<br>Reverse 5'-CCAGATAGATGTCTCCCAGGCTTG-3' | 350 | Present Study |
| *Myf5*$^{neo}$ | Forward 5’-CGTTGGCTACCCGTGATATT-3'<br>Reverse 5’-CAGCTCAGCTTTGTGTGCTC-3' | 670 | 1423602 |
| *Myf5*$^{loxP}$ | Forward 5'-GGTGTCTCCTCTCTGCTGAATCCAGGTAT-3'<br>Reverse 5'-AGGTGCACGCACGTGCTCCTCACTGTCTGA-3' | 349 | 15386014 |
| *Myf5* WT | Forward 5'-TGAAGGATGGACATGACGGAC-3'<br>Reverse 5'-CAGCTCAGCTTTGTGTGCTC-3' | 144 | Present Study |
| *MyoD*$^{iCre}$ | Forward 5'-GTCATTGTACTGTTGGGGTTCC-3'<br>Reverse 5'-AGCATCTTCCAGGTGTGTTCAGAG-3' | 254 | Present Study |
| *MyoD* WT | Forward 5'-GTCATTGTACTGTTGGGGTTCC-3'<br>Reverse 5'-CTTGAGCGTCTCGAAGGCCTC-3' | 485 | Present Study |
| *lacZ* | Forward 5’-CCGAAATCCCGAATCTCTATC-3'<br>Reverse 5’-TTGGCTTCATCCACCACATAC-3' | 333 | 10101129 |

**Supplemental Table 2. Coordinates of dREG candidate enhancers and regions cloned for transgenic analyses**

| | **Coordinates of Region(s) Selected by dREG** | | | **Coordinates of Regions Cloned for Transgenic Mice** | | |
|---|---|---|---|---|---|---|
| | Start | Stop | Size (bp) | Start | Stop | Size (bp) |
| -36lacZ | chr 7: 46,337,051 | chr 7: 46,339,152 | 2,102 | chr 7: 46,336,638 | chr 7: 46,342,463 | 5,826 |
| | chr 7: 46,341,501 | chr 7: 46,341,952 | 452 | | | |
| -60lacZ | N/A | N/A | N/A | chr 7: 46,315,210 | chr 7: 46,318,065 | 2,856 |
| -96lacZ | chr 7: 46,282,551 | chr 7: 46,285,802 | 3,252 | chr 7: 46,279,272 | chr 7: 46,279,272 | 7,081 |

**Supplemental Table 3.** BL-Hi-C and ChIP-seq datasets

| Dataset | Cell Type/Condition[1] | PMID | Data Availability | Accession Number(s) | Figure Number |
|---|---|---|---|---|---|
| **Hi-C Datasets** | | | | | |
| BL-Hi-C | Primary myoblasts/GM | 35017543 | CRA002490 | CRR126286, CRR126287, CRR126288, CRR126289, CRR126290, CRR126291 | 7A, Supp. 4A |
| BL-Hi-C | Primary myoblasts/DM | 35017543 | CRA002490 | CRR126292, CRR126293, CRR126294, CRR126295, CRR126296, CRR126297, CRR126298, CRR126299 | 7B, Supp. 4B |
| Hi-C | ESC | 29313530 | 4DNESU4BQU4G | 4DNFI3JYF9VS | Supp. 3 |
| **ChIP-Seq Datasets** | | | | | |
| H3K27ac | Primary myoblasts/GM | 35017543 | CRA002490 | CRR126346, CRR126347 | 7A, Supp. 4A |
| H3K4me1 | Primary myoblasts/GM | 35017543 | CRA002490 | CRR126354, CRR126355 | 7A, Supp. 4A |
| Histone input | Primary myoblasts/GM | 35017543 | CRA002490 | CRR126370 | 7A, Supp. 4A |
| CTCF | Primary myoblasts/GM | 35017543 | CRA002490 | CRR126330, CRR126331 | 7A, Supp. 4A |
| MYOD | Primary myoblasts/GM | 35017543 | CRA002490 | CRR126326, CRR126327 | 7A, Supp. 4A |
| TF input | Primary myoblasts/GM | 35017543 | CRA002490 | CRR126374 | 7A, Supp. 4A |
| H3K27ac | Primary myoblasts/DM | 35017543 | CRA002490 | CRR126348, CRR126349 | 7B, Supp. 4B |
| H3K4me1 | Primary myoblasts/DM | 35017543 | CRA002490 | CRR126356, CRR126357 | 7B, Supp. 4B |
| Histone input | Primary myoblasts/DM | 35017543 | CRA002490 | CRR126371 | 7B, Supp. 4B |
| CTCF | Primary myoblasts/DM | 35017543 | CRA002490 | CRR126332, CRR126333 | 7B, Supp. 4B |
| MYOD | Primary myoblasts/DM | 35017543 | CRA002490 | CRR126328, CRR126329 | 7B, Supp. 4B |
| TF input | Primary myoblasts/DM | 35017543 | CRA002490 | CRR126375 | 7B, Supp. 4B |
| H3K27ac | ESC | 32737473 | https://www.encodeproject.org | ENCFF583WVZ | Supp. 3 |
| H3K4me1 | ESC | 32737473 | https://www.encodeproject.org | ENCFF456IEO | Supp. 3 |
| CTCF | ESC | 22955616 | https://www.encodeproject.org | ENCFF144RYY | Supp. 3 |
| MYOD | C2C12/DM | 25409824 | GSE49847 | GSM915185 | 4 |
| MYF5 | Myf5-transfected MEFs | 26906734 | GSE75370 | GSM1954032 | 4 |
| MyoG | C2C12/DM | 25409824 | GSE49847 | GSM915163 | 4 |
| Mef2D-a2 | C2C12/DM | 23723416 | GSE43223 | GSM1058956 | 4 |
| H3K27ac | C2C12/DM | 28575289 | GSE93916 | GSM2464971 | 4 |
| DNase1 | Adult skeletal muscle | 25411453 | GSE51336 | GSM1014189 | 4 |

[1]All data are derived from mouse cells/tissue. GM, growth media; DM, differentiation media.
